# Dual-Lipa: A sequential multi-omic subcellular landscape map of mouse heart

**DOI:** 10.64898/2026.05.11.724441

**Authors:** Haoyun Fang, Alin Rai, Kevin Huynh, Seyed Sadegh Eslami, Thy Duong, Alex Faulkner, Peter Meikle, David W Greening

## Abstract

Spatial multi-omics can provide a unique understanding in molecular organisation and heterogeneity of organs including the heart. Heart function is dependent on the abundance and the spatial arrangement of proteins and lipids. Yet, an integrated multi-omics landscape of the heart at subcellular scale remains unknown. Here, we used a sequential lipid-proteomic analytical pipeline applied to the mouse heart to identify, map, and integrate lipid and protein features, revealing coordinated distinct molecular networks within defined subcellular niches. Here, we benchmark this dual extraction workflow for integrated global proteome-lipidome analyses to demonstrate its performance. We developed conserved subcellular proteome of mouse heart across 14 niches and applied the knowledge of subcellular proteome to spatially resolve the heart lipidome, identifying unique lipid features from different subcellular niches, including mitochondria (e.g., cardiolipin, LPC) and plasma membrane-enriched lipids (e.g., plasmalogens). We identified sex-dependent molecular differences in heart subcellular proteome between male and female, including RNA-protein complexes, mitochondrial calcium handling and immune regulatory pathways. We demonstrate that lipid-protein integrated multi-omics analysis of the heart by dual extraction workflow and mass spectrometry could enable previously unidentified discoveries in heart molecular composition and organisation and spatial cardiac biology.

**Highlights:**

- Resolved tissue-wide subcellular architecture of cardiac tissue through simultaneous high-resolution proteomics and lipidomics profiling.
- Systematically evaluated MS-compatible dual extraction strategies for integrated proteome-lipidome recovery and performance.
- Proteomics revealed major sex-dependent differences in mouse left ventricles, primarily involving cellular metabolism, immune regulation, detoxification pathways, and signalling responses.
- Lipidomics identified strong sex-specific lipid signatures, including differences in omega-3 vs omega-6 fatty-acid containing lipid species, triacylglycerides, and acylcarnitines.
- Developed supervised machine-learning models to classify 14 subcellular niches from tissue-wide fractionation proteomics and identified a conserved core subcellular proteome.
- Applied the tissue-wide subcellular proteome model to infer lipid subcellular localisation, revealing mitochondria-enriched lipids (e.g., cardiolipin, LPC) and plasma membrane-enriched lipids (e.g., plasmalogens) in the mouse left ventricle.
- Mapped sex-specific subcellular distribution in the heart, highlighting pathways linked to RNA-protein complexes, mitochondrial calcium handling, and immune regulatory networks.

## Introduction

The heart is a highly structured and metabolically active organ whose function depends on coordinated cellular interactions, precise molecular regulation and communication between specialised organelles^1–3^. The regulation of organelle composition and dynamics is important for supporting cardiac functions including calcium signalling^4–6^, excitation-contraction coupling^7–10^, cellular energetics^11–14^, redox balance^15–19^, and proteostasis^19–22^. Dysregulated organellar composition and impaired inter-organelle communication are frequently reported in pathological conditions such as myocardial infarction^23,24^, arrhythmia^25,26^ and heart failure^27–31^. Apart from cardiac pathological remodelling, subcellular organisation is also important for understanding physiological cardiac differences, including biological sex - a major biological factor that shapes differential cardiac molecular regulation.

Due to the differences in sex hormones and chromosome, male and female hearts differ in cardioprotective capacity and susceptibility to different cardiovascular diseases^32^. Recently, omics-based cardiovascular studies revealed sex-dependent molecular signatures in cellular metabolism, immune regulation and cellular detoxification^33–36^. Growing evidence also suggests that sex-dependent cardiac differences may extend to subcellular organelle composition and organisation, where functional differences were reported in mitochondrial biogenetics and resilience, sarcoplasmic reticulum calcium transients and magnitude of contractile apparatus shortening between male versus female hearts^37,38^. To date, most sex-based transcriptomic, proteomic and lipidomic studies of the heart have been performed at the bulk tissue level^35,39–41^, leaving the spatial distribution of biomolecules across subcellular compartments rarely investigated. This suggests a need to investigate sex-dependent cardiac molecular organisation at subcellular resolution.

Proteins and lipids are central molecular components of cardiac subcellular organisation. Here, proteins regulate enzymatic activity, signalling dynamics and structural organisation, while lipids define organelle membrane architecture, cellular energetics and specialised signalling and cellular processes^42–45^. Importantly, the functional roles of these biomolecules are not only driven by their abundance, but also highly dependent on their subcellular distribution and spatial dynamics to preserve organelle structure and function in homeostasis^43,46^. At the organelle level, protein and lipid dynamics together help maintain local membrane environment, organelle morphology, inter-organelle communication and compartment-specific function^47–49^. Therefore, integrated characterisation of proteins and lipids is required to understand cardiac molecular subcellular organisation and how it varies across physiological contexts such as biological sex.

Recent advances in correlation profiling-based subcellular proteomics have enabled cell-wide and tissue-wide mapping of protein localisation across distinct subcellular niches. Experimental and computational frameworks such as LOPIT^50,51^, DOM^52,53^ and SubCellBarcode^54^ perform partial subcellular fractionation of cells or tissues to generate proteomic profiles across resolved fraction gradients. Using curated organelle-specific marker proteins, these pipelines enable the simultaneous mapping of multiple subcellular niches through machine learning-based classification of protein co-fractionation patterns. These approaches have been extended to investigate spatial regulation of protein post-translational modifications^55,56^, newly synthesised proteins^57^ and to integrate additional layers of omics data^58,59^, including defining proteome features and subcellular organisation of the heart^60^.

Despite these advances, tissue-wide mapping of the subcellular lipidome remains challenging. It is because most lipids are shared across multiple organelles and the existence of organelle-specific lipids are limited^46^. In addition, the subcellular lipid distribution is highly dynamic from vesicle trafficking, membrane remodelling and transient organelle contact sites^61,62^ which further complicate the direct assignment of lipid species to subcellular niches. Recently, computational frameworks using spatial proteomic information as a reference have demonstrated the feasibility of predicting lipid localisation from proteome-wide fractionation patterns^59^. This provides a strategy to reconstruct lipid distributions across subcellular compartments and to integrate lipidomic and proteomic spatial information. However, integrated workflows that combine lipid-protein extraction, high-sensitivity mass spectrometry, subcellular fractionation and computational spatial modelling remain limited in cardiac tissue, particularly for investigating physiological variables such as biological sex.

We introduce Dual LiPA, an integrated lipidome-proteome analytical framework for investigating the spatial molecular organisation of the mouse heart, particularly focusing on left ventricle. We first evaluated the performance of different sequential lipid-protein extraction and analytical workflows using MS-based proteomic and lipidomic data of bulk cardiac tissue. We then conducted differential centrifugation-based subcellular fractionation followed by concurrent proteomics and lipidomics analysis. For the classification of subcellular proteins, we applied a binary relevance strategy with random forest-based multi-label classification to define a conserved cardiac subcellular proteome comprising more than 3,000 proteins across 14 subcellular niches. Building upon this spatial proteome map, we reconstructed subcellular lipid distributions using protein localisation as spatial anchors and uncovered organelle-associated lipids from the heart. By integrating proteomic and lipidomic data layers, we highlight the coordinated lipid-protein organisation across membrane-rich subcellular niches of left ventricle. Our analysis revealed a sex-dependent subcellular proteome localisation patterns different between male and female sex across organelle compartments, associated with transcriptional regulation, immune responses and ion homeostasis. This framework enables an integrated spatial map of lipid-protein organisation of the heart, and refined understanding in biological molecular patterns of lipids and proteins across distinct organelles under biological conditions.

## Experimental procedures

### Mouse heart tissue source

All mouse experiments and tissue collection were approved by the Alfred Research Alliance Animal Ethics Committee (Victoria, Australia; approval number P2580). Wild-type female and male C57BL/6 mice (8-10 weeks old) were sourced from AMREP AS Pty Ltd (VIC), housed in AMREP AS Pty Ltd facility with 12:12 h light-dark cycle and fed with standard mouse chow. Following euthanasia, hearts were isolated immediately, rinsed in ice-cold PBS, left ventricle (LV) dissected, and snap-frozen on dry ice.

### Tissue homogenisation and subcellular fractionation

Tissue homogenisation and subcellular fractionation were conducted as described^60^. In brief, around 30 mg of LV tissue were transferred into 1.5 mL safe lock microtube with prechilled 2.0 mm zirconium oxide beads and 1 mL of hypotonic buffer (0.25 M sucrose, 10 mM HEPES pH 7.4, 2 mM EDTA, 2 mM magnesium acetate tetrahydrate, 200 µM butylated hydroxytoluene) with Halt Protease and Phosphatase Inhibitor Cocktail using a pre-chilled Bullet Blender (Next Advance) with two rounds of homogenisation (setting 10 for 15 s, setting 3 for 15 s) followed by 10 mins of gentle rotation at 4°C. Homogenates were clarified at 100*g* for 5 min in a fixed-angle rotor centrifuge (Eppendorf 5804R; TLA-55 rotor). A 500 μl aliquot of the lysate was reserved for total heart (LV) proteome and lipidome analysis.

Tissue homogenates were sub-aliquoted for (i) global tissue profiling (15 μl) or (ii) differential fractionation (485 μl). For the latter, samples were centrifuged three times at 200*g* for 5 mins with supernatant collected for differential centrifugation-based subcellular fractionation using Eppendorf 5804R or Optima MAX-XP with fixed-angle TLA-55 rotor (Beckman Coulter) at 4 °C. For heart organelle differential fractionation, a 10-step differential centrifugation workflow was employed as previously described^60^. The pellets from each centrifugation step (F01-F10) collected and reconstituted in 30 µL hypotonic buffer. Supernatants following the F10 were collected as F11. Protein concentrations (global, F01-F11) were quantified using the microBCA assay (Thermo Fisher, 23235).

For global multi-omic analysis, proteins from the same biological replicates were sub-aliquoted to four tubes (2 µg) for sequential lipid extraction and proteomic analysis. For subcellular multi-omic analysis, normalised fractions (2 µg of protein) were assigned for sequential lipid extraction and proteomic analysis. All samples were lyophilised and stored in −80 °C for subsequent extraction and analysis.

### Single-phase lipid extractions

For global heart extract, we applied different lipid extraction procedures, including different butanol-and-methanol-based approaches (BUME, Hydro-BUME) and acetone-based extraction (Acetone). For BUME, lipid extraction was conducted as described^63^, where lyophilised samples were reconstituted in 10 µL of LC-MS grade water with 100 µL BUME solvent (butanol:methanol at 1:1 ratio) supplied with lipid internal standards (Supp Table 1). Samples were extensively vortexed followed by 1 hr water bath sonication at RT. Samples were centrifuged at 13,000 g for 10 mins at RT, with supernatants transferred into glass vials and stored in −80 °C for subsequent targeted lipidomic MS acquisition. The remaining pellets were dried (SpeedVac) and stored in −80 °C for downstream proteomic analysis. For Hydro-BUME, we modified solvent ratio (butanol:methanol:water at 2:2:1 ratio with 50 µM medronic acid and 1 mM ammonium formate). For heart subcellular fraction analysis, lipid extractions were conducted with Hydro-BUME approach. For Acetone approach, we replaced BUME with Acetone solvent (88% Acetone in water) and collected the supernatant following centrifugation at 13,000 g for 10 mins at RT. The fraction was lyophilised and reconstituted in 1:1 butanol:methanol with sonication in a water bath for 15 mins before transfer into glass vial inserts.

### Proteomic sample preparation

A total of 50 μg of protein was adjusted (2% sodium deoxycholate and 100 mM Tris-HCl, pH 8.5) and boiled for 5 min at 95 °C and 1,000 rpm. Protein pellets obtained from each lipid extraction workflow (BUME, Hydro-BUME, Acetone, None) were applied. Pellets post lipid extractions were subjected to bead-based SP3 protein workflow as described (19, 78). Proteins were lysed in SDS/HEPES buffer (50 mM HEPES, pH 8.0) supplemented with HALT protease/phosphatase inhibitors, then denatured in 2% SDS and reduced with 10 mM DTT followed by alkylation with 20 mM iodoacetamide (IAA) in the dark. Excess IAA was quenched by adding DTT. Proteins were subsequently captured on Sera-Mag magnetic beads (SP3), bound in the presence of ethanol, and washed repeatedly with 80% ethanol to remove detergents and other contaminants. Bead-bound proteins were then digested for 6 hr at 37 °C using sequence-grade trypsin and Lys-C (enzyme-to-protein ratios of 1:50 and 1:100, respectively). Peptide digests were acidified to 2% formic acid, and lyophilised and reconstituted in 10 µL 0.7% (v/v) trifluoroacetic acid in LC-MS grade water.

### Lipidomic data acquisition

LC-MS method was applied as previously described^64^. In brief, the lipid extracts were analysed on Agilent 1290 series HPLC system coupled with Agilent 6495C QQQ mass spectrometer. The lipid extracts were separated on a ZORBAX eclipse plus C18 column (2.1□× 100□mm 1.8□µm, Agilent) with using solvent A (50:30:20, water: acetonitrile: isopropanol) and solvent B (1:9:90, water: acetonitrile: isopropanol) at flow rate of 0.4 mL/min with column temperature at 45°c. The chromatography settings are as follows: Starting at 15% B, going to 50% at 2.5 minutes, 57% at 2.6 minutes, 70% at 9 minutes, 93% at 9.1 minutes, 96% at 11 minutes, 100% at 11.1 minutes and holding until 12 minutes before going down to 15% at 12.2 minutes and re-equilibrating at 15% B until 16 minutes. The mass spectrometer settings are as follows: gas temperature, 200°c, gas flow, 17l/min, nebuliser pressure, 20psi, sheath gas temperature, 280°C, sheath gas flow, 10l/min, capillary voltage, 3500V, nozzle voltage, 1000V. Positive and low-pressure RF was set to 210 and 160 respectively. Isolation widths for Q1 and Q3 were set to “unit” resolution at 0.7 amu. The detailed mass spectrometry settings and transitions for each lipid class were recorded in Supp Table 2.

### Lipidomic data integration

Raw lipidomic MS data was processed using MassHunter Quant 10.0 (Agilent Technologies). To generate relative lipid concentration data per sample, the peak areas of different lipid species were normalised to their corresponding lipid internal standards. Potential isomeric lipid species without structural resolution were annotated with (a), (b) or (c) suffixes to highlight different elusion orders. Lipid class totals were calculated by summing up individual lipid species belonging to the same lipid class.

### Proteomic data acquisition

Peptide samples were analysed using Thermo Fisher Neo UHPLC system coupled with Orbitrap Astral mass spectrometer in data-independent acquisition mode. One microlitre (100 ng equivalent) peptide was injected from all samples.

For left ventricle bulk tissue proteome samples, peptides were loaded to PepMap™ Neo Trap Cartridge (#174500, Thermo Fisher Scientific) via trap-and-elute mode, and separated on an EASY-Spray PepMap RSLC C18 column (#ES906; Thermo Fisher Scientific) at 55 °C in proteomics solvent A (0.1% LC-MS grade formic acid in LC-MS grade water) and proteomics solvent B (0.1% LC-MS grade formic acid in 80% LC-MS grade acetonitrile prepared in LC-MS-grade water). Peptides separation was conducted with a gradient of different percentage of Solvent B starting with 4% hold for 0.7 min, 4-8% over 0.3 min, 8-8.1% over 0.1 min, 8.1-28% over 8.7 min, 28-50% over 3.5 min, 50-99% over 0.5 min and held at 99% for 0.7 min. The flow rates change from 3 µL/min to 0.9 µL/min at 1.1 min, 0.9 uL/min to 2.5 µL/min at 13.8 min. Full MS scans were acquired with orbitrap mass analyser at 240,000 resolution over 380-980 m/z with AGC target at 5e6, maximum injection time at 5 ms with 1 microscan. MS/MS spectra were acquired with Astral analyser with 200 isolation windows of 380-980 m/z range with 0 m/z overlap with the same precursor range. Further MS/MS parameters were set to AGC target at 5e4, maximum injection time at 5 ms, 1 microscan, HCD normalised collision energy at 25%, fragment ions range of 150-2000 m/z and total cycle time of 0.6s.

For subcellular fraction samples, peptides were loaded to trap column (Acclaim PepMap100 C18 3 μm beads with 100 Å pore-size, Thermo Fisher Scientific) via heated trap-and-elute mode at 55 °C and separated by analytical column (1.9-μm particle size C18, 0.075 × 250 mm, Nikkyo Technos Co. Ltd) with external heating (butterfly portfolio heater, Phoenix S&T) at 55 °C. Peptides separation was conducted with a gradient with different percentages of solvent B. Peptides separation was conducted with a gradient with different percentage of Solvent B starting at 4%, 4-10% over 1 min, 10-50% over 8 min, 50-99% over 2 min and held at 99% for 3 min. The flow was kept at 600 µL/min. Full MS scans were acquired with orbitrap mass analyser at 240,000 resolution over 380-980 m/z with AGC target at 5e6, maximum injection time at 5 ms with 1 microscan. MS/MS spectra were acquired with Astral analyser with 300 isolation windows of 380-980 m/z range with 0 m/z overlap with the same precursor range. MS/MS parameters were set to AGC target at 5e4, maximum injection time at 3 ms, 1 microscan, HCD normalised collision energy at 25%, fragment ions range of 150-2000 m/z and total cycle time of 0.6 s. MS spectra were acquired using Xcalibur software v4.7 SP1 (Thermo Fisher Scientific).

### Proteomics data processing

DIA raw data files were processed using DIA-NN (v2.0) in the “robust LC (high precision)” mode with retention time (RT)-dependent normalization enabled, searching against the UniProt databases for mice (UP000000589_10090, 54,791entries)^65^. Search parameters included library-free mode with selected “FASTA digest for library-free search/library generation”, “Deep learning-based spectra, RTs and IMs prediction” and enabled matching between runs function. Trypsin/P with maximum 1 missed cleavage was selected for enzymatic digestion. The precursor charge range was set to 1-4 with range between 300-1800 m/z. Peptides search parameters set to 7-30 amino acids in length. N-term methionine excision and cysteine carbamidomethylation set as fixed modification and variable (no variable modification) modifications as default. Mass accuracies at MS1 and MS2 were automatically determined by DIA-NN via default setting (Mass accuracy: 0.0). Mass spectra were analysed using default settings with a false discovery rate (FDR) of 1% for precursor identifications. Machine learning was set to “NNs (cross-validated)” and quantification strategy set to “Quant UMS (high precision).

### Statistical analysis for global proteomic and lipidomic data

Global proteomic and lipidomic analyses were conducted using R (4.5.0). For lipidomic analysis, the raw lipid intensities were used to assess lipid recovery for different extraction workflows. We further applied lipid relative abundance normalised to lipid internal standards (Supp Table 3), with log2 transformation. For differential analysis of lipid species, a linear mixed-effect model was fitted using lmerTest package^66^. The model estimates the fixed effect of biological variance (i.e., male and female sex dependent biological effect) and extraction workflow (i.e., Acetone, BUME, Hydro-BUME) while accounting for repeated measurements from the same biological replicate via a random intercept.

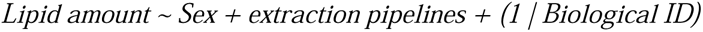

The estimates, T-statistics, p-values, and Benjamini-Hochberg adjusted p-values of the fixed effects (Sex, Pipelines) are reported. Lipid annotation was constructed in-house to describe the lipid category, lipid class, lipid subclass and features of all quantified lipid species (Supp Table 4). Lipid set enrichment analysis was conducted using the in-house lipid annotation and GSEA() function from clusterProfiler package^67^.

For proteomics data analysis, data quality cut-off was applied to only include protein groups with more than 1 peptide and quantified in at least 75% in at least one group (i.e., 3 out of 4). Protein group intensities were log2 transformed, with missing values imputed using msImpute() function with group-specific settings and “v2-mnar” mode from msImpute package^68^. Differential analysis was assessed using limma linear modelling framework using limma package^69^. Biological and analytical variance (i.e., male and female sex dependent biological effect, extraction workflow) were set as factors with sum-to-zero contrast. To account for repeated measurements of the same biological samples, duplicateCorrelation was enabled for individual biological samples. Empirical Bayes moderation (eBayes) was applied, and moderated statistics (log2 fold change, t-value, p-value, FDR) were extracted for each model coefficient using topTable(). Gene ontology-base and reactome-based functional enrichment analysis was conducted using GSEA() and compareCluster() function from clusterProfiler package.

### Subcellular proteomics and lipidomic data processing and machine learning

Subcellular proteomics and lipidomics data processing were conducted in R using limma, dplyr, and msImpute packages. For subcellular proteome data, protein groups identified with 100% in at least one group (identified in all n = 3) were subjected to downstream process. Protein group label-free quantification (LFQ) intensities were log2-transformed and normalised via cyclic loess at group-specific level using normalizeCyclicLoess() function from limma package. Missing values were imputed at group-specific level using msImpute() function with “v2-mnar” setting from msImpute package. For subcellular lipidomics data, lipid-internal standard normalised lipid abundance data was used (Supp Table 5-6). Normalised and imputed protein and lipid data were back transformed (2^x), and these values were scaled to 0-1 sum intensity ratio across 11 subcellular fractions of each biological sample, namely fraction profiles.

To curate subcellular protein markers for machine learning-based assignment, we initially assigned subcellular protein marker candidates based on GO cellular compartment annotation^70^ and other subcellular markers curated from previous studies^50,52,60^. We further compared fractional profiles of all protein groups within/against each group (biological replicates), with stringent inclusion criteria of protein groups with >0.9 Cosine correlation for pairwise comparisons between all biological replicate pairs included. Further, we ensured protein markers are dependent on their biological difference, where Wilcoxon t-test of Euclidean distance of protein fraction profiles “between groups” (biological variance of female vs male) and “within groups” (male vs male, female vs female) were performed. Proteins (p<0.05) in “within group” distance less than “between group” distance were excluded for subcellular organelle protein marker selections.

For machine learning model construction, binary relevance approach was adapted for multi-label classification of proteins and lipids to different subcellular niches. In detail, the subcellular protein map was converted to binary label matrix where protein markers were annotated as 1 for the destinated subcellular niche and 0 for remaining subcellular niches. Using caret package in R^71^, independent random forest (RF) models were trained on each subcellular niches using the protein fractionation intensity profiles (F01-F11) as input features. For hyperparameter optimisation, each model was tuned using grid search of mtry (1-16) and evaluated with 5 repeats of 10-fold cross-validation. F1-score of precision-recall statistics determined on the optimal mtry for each RF model. Class probability was enabled for output data. Following training, each RF model was used to predict the probability across all subcellular niches based on subcellular proteomics and lipidomics data, where class probability of protein and lipid assignment to each subcellular niche was assembled into a matrix.

### Statistics for comparison of subcellular localisation

Differential protein fraction profiles were assessed using generalised additive model (GAM) via mgcv package^72^. For each protein, 11 subcellular fractions were assigned numeric covariate modelled by intensity ratios with biological replicates accounted as random effect. Overall, two GAMs were fitted for each protein. The null hypothesis model was constructed based on shared subcellular fractionation profiles independent of biological group (female, male), whereas the alternative model accounted group-dependent differences which assigned group as a categorical factor. Likelihood ratio testing was conducted between null model and alternative model base on chi-square distribution. Valid p-values were extracted and BH-adjusted for multiple testing. To evaluate the magnitude of subcellular profile differences between groups, pairwise cosine distances (1-cosine correlation) of subcellular fractionation profiles between groups were calculated, and the mean cosine distance reported.

### Bioinformatic analysis and data visualisation

For proteomic data, GO- and KEGG-based gene set enrichment analysis and overrepresentation analysis were conducted using clusterProfiler in R. For lipidomic data, lipid set enrichment analysis and overrepresentation analysis were conducted using clusterProfiler, incorporating an in-house constructed lipid annotation table (Supp Table 4) comprising lipid domains, classes, subclasses and features (fatty acyl chain length and saturation). For data visualisation in bar plots, volcano plots, boxplots, line plots scatter plot and lollipop plots, ggplot2^73^was used. UMAP segregation of subcellular proteome from individual biological replicates were conducted using uwot package^74^ with 500 n_neighbours and 0.5 min_dist. Subcellular lipidome data were projected to subcellular proteome using the UMAP model established with proteome data. UMAP-informed protein and lipid clustering were visualised in using ggplot2 package. Gseaplot, cnetplot and emaplot of gene set and lipid set enrichment analysis are visualised using enrichplot package^75^.

### Data availability

Proteomics and lipidomics data generated in this study (raw files, processed searchable data) are available in the ProteomeXchange Consortium via the MassIVE partner repository and available via MassIVE with identifier (MSV000101429). RF-based multi-label model and the performance output are accessible via GitHub repository^76^.

## Result

### Single-phase lipid extraction enables concurrent lipidomic and proteomic profiling from limited biological samples

To achieve heart-centric lipidome and proteome co-profiling, we evaluated experimental workflows to extract, correlate, and analyse the lipid and protein features simultaneously from male and female left ventricles (Fig 1a). Briefly, native dissected left ventricle region was processed for single-phase lipid extraction, in combination with SP3-based proteomic sample preparation workflow (Fig 1a). From limited (3 µg protein-equivalent) heart extract, we compared extraction efficiency and intensity/coverage of different aqueous monophasic extractions including Hydro-BUME designed to improve recovery of phosphorylated lipid species and polar amphiphilic lipids, butanol–methanol extraction (BUME)^63^, and acetone-based (Acetone) protein extraction, used in proteomic workflows for delipidatation as a pseudo-lipid extraction approach^77^.

**Figure 1.**
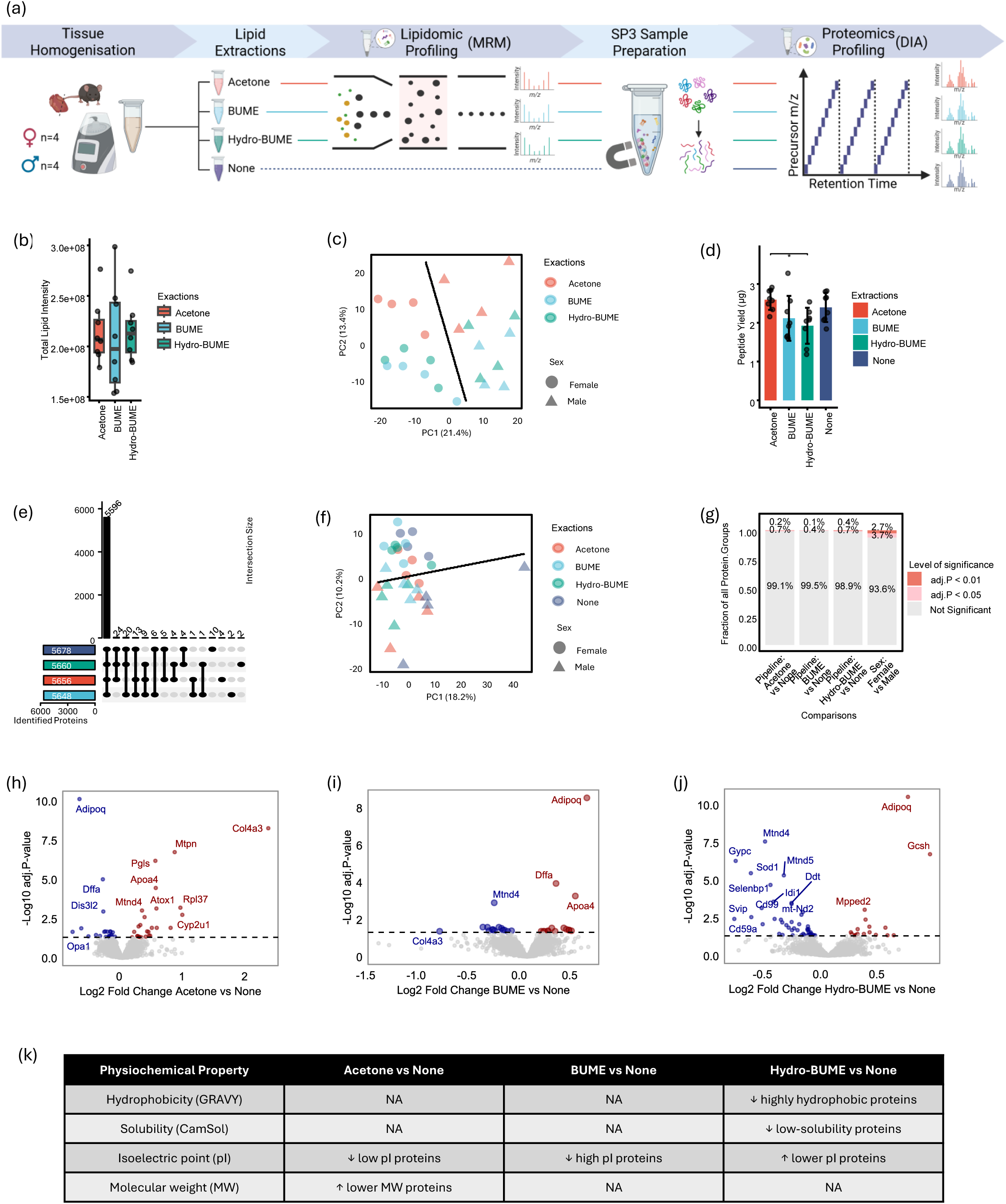
Evaluation of a dual lipid–protein sample preparation workflow for integrated MS-based lipidomic and proteomic profiling.(a) Experimental workflow for simultaneous lipidomic and proteomic profiling. Left ventricles from female and male mice were collected and either processed directly (None) or subjected to lipid extraction using different solvent systems (Acetone, BUME, Hydro-BUME). Lipid extracts from the three extraction methods were analysed by targeted MRM DIA LC–MS/MS lipidomics, while the remaining protein pellets were retained for proteomic analysis. Protein extracts, including those without prior lipid extraction, were processed using SP3-based sample preparation and analysed by DIA LC–MS/MS.(b) Boxplots showing the summed lipid intensities obtained from Acetone, BUME and Hydro-BUME extraction-derived lipidomic data. Each dot represents one biological sample. (c) PCA of quantified lipidomic profiles of different biological sex and sample preparation pipeline. Linear discriminant analysis (LDA) based on PC1 and PC2 scores illustrates predicted sex separation across PCA space.(d) Peptide yield recovered from different sample preparation pipelines. Peptide yield was analysed using repeated-measures ANOVA with biological sample as the repeated measure, followed by Tukey-adjusted post hoc comparisons. Bars show mean ± s.d.; points represent individual samples; asterisks indicate adjusted P values (Padj < 0.05).(e) UpSet plot showing commonly and differentially identified protein groups across different sample preparation pipelines.(f) PCA of log_₂_-transformed proteomic intensities from different biological sex and sample preparation pipeline. LDA based on PC1 and PC2 scores illustrates predicted sex separation across PCA space.(g) Stacked bars showing the fraction of protein groups significantly associated with sex or extraction pipeline (adjusted P < 0.05 or < 0.01) based on empirical Bayes–moderated statistics (limma) accounting for repeated measurements from the same biological sample. P values were adjusted using the Benjamini–Hochberg correction.(h–j) Volcano plots showing differential log2-transformed protein LFQ intensities for each lipid extraction workflow (Acetone (h), BUME (i), Hydro-BUME (j)) compared with the baseline condition (None). Differential analysis was performed using limma accounting for repeated measurements from the same biological sample. P values were adjusted using the Benjamini–Hochberg correction. Proteins with significantly higher or lower intensities (Padj < 0.05) are highlighted in red and blue, respectively.(k) Summary of physicochemical properties of proteins significantly different in each lipid extraction workflow relative to the baseline condition (None).

We demonstrate all extractions yield similar level of total lipid intensity, measuring 779 lipid species across all extractions (Fig 1b). Hydro-BUME showed slightly higher total lipid intensity and slight elevated reproducibility. At lipid class level, there is minimal lipid class-wide bias between different lipid extraction strategies (Supp Fig1a, Supp Table 6). Free cholesterol (COH), lysoalkylphosphatidylcholine (LPO O) and oxidised lysophosphatidylcholine (oxLPC) lipids presented with higher intensity by Acetone extraction, while intensities of cardiolipins (CL), oxidised alkylphosphatidylcholine (oxPC O) and phosphatidylinositol monophosphate (PIP1) are elevated in BUME and highest in Hydro-BUME extraction, suggesting Hydro-BUME-based extraction represents a promising alternative to extract a diversity of biological lipid types (Supp Fig 1a). We evaluated lipidomic quantitative variability by comparing coefficient of variation (CV) of internal-standard-normalised lipid abundances of every lipid species across three different extraction processes (Supp Fig1b, Supp Table 6). The three extraction workflows all share similar distribution of CV with Hydro-BUME has the lowest median CV (25.5%) comparingly to BUME (26.1%) and Acetone (26.6%) (Supp Table 6). PCA analysis showed that Acetone-derived lipidomes clustered further away from the BUME and Hydro-BUME groups, suggesting that acetone extraction produced a slightly more distinct lipidomic profile. Notably, biological sex differences remained the dominant source of variance (PC1: 21.4%) (Fig. 1c), indicating extraction effects did not mask the true biological lipidomic variations between sex.

Next, we acquired concurrent proteomic profiling of each lipid extraction workflow from heart, relative to non-lipid extraction proteome (None) as baseline controls (Fig. 1a). While BUME detected the least number of quantified peptides relative to non-lipid proteome workflow (Supp Fig. 1c), the protein-level coverage exhibited high consistency across all lipid extraction workflows (Supp Fig1d, Supp Table 8). We found high proportion of protein groups (98%, 5,596 proteins) commonly identified in each extraction workflow (Fig. 1e). Notably, PCA showed that proteomes generated from different extraction methods were highly similar, indicating that the extraction workflow did not introduce major technical variation or mask the underlying biological separation between male and female heart proteomes along (Fig. 1f).

To evaluate proteome differences introduced by lipid extraction, we compared extraction-derived proteomes with the baseline proteome (None) and found that only around 1% of proteins showed significantly altered intensity (adjusted p < 0.05; Acetone: 0.9%; BUME: 0.5%; Hydro-BUME: 1.1%; Fig. 1g–j, Supp Table 9). We then evaluated the physicochemical properties of significantly altered proteins from each comparison, focusing on hydrophobicity (GRAVY), predicted solubility (CamSol), isoelectric point (pI) and molecular weight (MW). In comparison to the baseline proteome, each workflow showed mild physicochemical bias. Acetone workflow favoured recovery of low-MW proteins and reduced proteins with low pI (Supp. Fig. 2a, Fig. 1k, Supp Table 9). BUME reduced recovery of high-pI proteins (Supp. Fig. 2b, Fig. 1k). Hydro-BUME reduced proteins with high hydrophobicity, low solubility and high pI while enriching lower-pI proteins, potentially reflecting the acidic extraction solvent (Supp. Fig. 2c, Fig. 1k). Together, this suggests that concurrent lipidomic and proteomic profiling of cardiac tissue is feasible, but proteome recovery remains partly influenced by extraction chemistry.

Despite physiochemical differences in proteome coverage based on lipid extraction workflow, it is important to highlight that lipid extraction-associated variation (∼1%) remained substantially limited relative to biologically driven proteome differences between different sex (∼7%) (Fig1g). This therefore enables our sequential workflow to identify lipid and protein features in a complex, biologically heterogeneous sample limited sample such as the heart.

### Applying dual lipid-protein workflow to decipher sex difference in heart

After evaluating the technical performance of dual lipid-protein analysis workflows from heart, we next assessed each lipid extraction approach dependent on accurately identify differences associated in the context of biological sex (Supp Fig 3). It is achieved by performing quantitative proteomics on different heart samples from healthy male and female mice following differential lipid extraction.

We show at a proteome level, 242 protein groups significantly higher (P.adj < 0.05) in female and 102 significantly higher (P.adj < 0.05) in male (Fig 2a). Among these proteins, 2 Y-chromosome expressing proteins in male (Ddx3y, Eif2s3y) and 12 X-chromosome expressing proteins in female (Fig2a, Supp Table 9), of which 9 (Cfp, Ddx3x, Eif2s3x, Hmgn5, Igbp1, Mpp1, Pdk3, Septin6, Uba1) have been previously reported as X-chromosome escapee proteins^41^. To understand functional difference at a proteome level of the heart across sex, we conducted KEGG-based over representation analysis on the sex-enriched protein groups, revealing female association with detoxication processes (“metabolism of xenobiotics by cytochrome P450”and “Glutathione metabolism”), “ferroptosis”, carbohydrate metabolism (“pentose and glucuronate interconversions”, “pyruvate metabolism”, “biosynthesis of various nucleotide sugars” 4). “nucleocytoplasmic transport”, and Akt-related signalling pathways (“adipocytokine signalling pathway, “ErbB signalling pathway”) (Fig 2b, Supp Table 10). Gene Ontology enrichment analysis further highlight enrichment of “protein folding chaperone”, “endocytic vesicle” and “nitric oxide metabolic process” in female proteome (Fig 2c, Supp Table 11). In contrast, male hearts are enriched with KEGG terms associated with “complement and coagulation cascades”, “cytoskeleton in muscle cells”, “fatty acid biosynthesis” and GO terms including “mitochondrion organization” and “hormone metabolic process” (Fig 2b-c). Despite distinct enrichment profiles, several biological pathways were shared between sexes but driven by different protein components (Fig. 2c), including complement activation, antioxidant response including glutathione conjugation and detoxification while male proteome group reflect enrichment of limiting membrane lipid peroxidation. Overall, these proteomic patterns showed strong concordance with previously reported sex-dependent molecular signatures ^35^.

**Figure 2.**
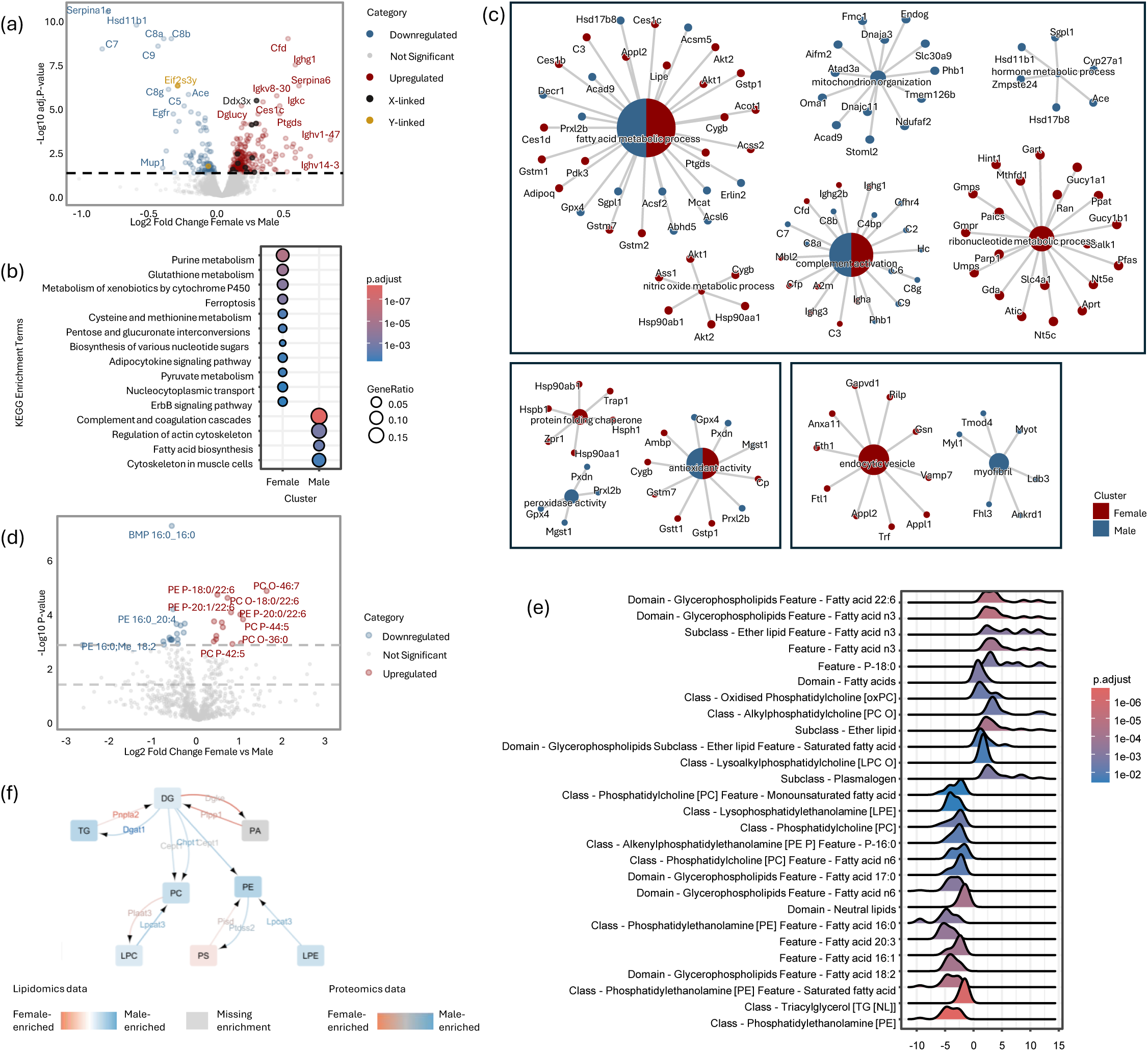
Sex-dependent differences in the proteome and lipidome of mouse left ventricles. (a). Volcano plots showing differential log2-transformed protein LFQ intensities of female versus male left ventricle proteome. Differential analysis was performed using limma accounting for repeated measurements from the same biological sample. P values were adjusted using the Benjamini–Hochberg correction. Proteins with significantly female or male enrichment (Padj < 0.05) are highlighted in red and blue, respectively. X-linked (black) and Y-linked (yellow) proteins are also highlighted. (b). Dot plot of KEGG pathways significantly enriched in female and male hearts based on over-representation analysis of proteins significantly enriched in each sex. (c). Protein-term network plots of significantly enriched GO terms (GOBP, GOCC and GOMF) and their associated differentially expressed proteins from female (red) and male (blue) enriched left ventricle proteomic data. (d). Volcano plot showing differential lipid species abundances (pmol/ug protein) between female and male left ventricle samples. Differential analysis was performed using linear mixed-effects modelling accounting for repeated measurements from the same biological sample. P values were adjusted using the Benjamini–Hochberg correction. Lipids with significantly female or male enrichment (Padj < 0.05) are highlighted in red and blue, respectively. (e). Ridge plot showing lipid set enrichment analysis (LSEA) of curated lipid annotation terms that are significantly (Padj < 0.05) associated with sex. (f). Integrated lipid–protein metabolic network derived from lipidomic profiles mapped to predicted lipid metabolic reactions using the Biopan platform and LIPIDMAPS enzyme annotations, visualised in Cytoscape. Node colours indicate lipid fold changes between female and male hearts, and edge colours indicate fold changes of the associated metabolic enzymes.

Lipidomics identified sex-specific lipid signatures, including 14 lipid species higher in female heart and 17 significantly elevated in male (Fig 2d, Supp Table 12). Lipid set fgsea-based enrichment analysis revealed ether-linked, oxidized and omega-3 fatty acyl-containing glycerophospholipids in female heart lipidome, while saturated and monounsaturated and omega-6 fatty acyl-containing glycerophospholipids and various neutral lipids, including triacylglycerols enriched in male (Fig. 2e, Supp Fig4a, Supp Table 13). Lipid species like proteome analysis in female hearts showed higher abundance of lipolysis-associated proteins, while for male heart proteome enriched in networks linked to lipid synthesis and storage pathways. Enrichment of ether lipids in female hearts is consistent with the established antioxidant role of ether lipids, particularly plasmalogens, which function as scavengers of reactive oxygen species^78–80^. This observation agrees with our proteomic findings showing enrichment of antioxidant-associated proteins in female hearts, collectively indicating coordinated molecular features associated with enhanced handling of oxidative stress.

Next, we mapped lipidomic profiles to predicted lipid metabolic reactions using the Biopan platform^81^ and enzyme annotations from LIPIDMAPS proteome database^82^ to reveal concordant multi-omics patterns across lipid and proteome level (Fig. 2f). We found triacylglycerol biosynthesis enzyme Dgat1 higher in males, while TG hydrolysis enzyme Pnpla2 higher in females, associated with similar lipidomic enrichment of TG species in male hearts. Similarly, phospholipid biosynthesis enzyme Lpcat3 was elevated in males, while phospholipid hydrolysis enzyme Plaat3 higher in females, consistent with observed the higher PC and PE lipid abundance in male.

Therefore, this dual lipid-proteomic workflow can profile lipidome and proteome from limited, biologically complex samples without biasing biological data quality dependent on the extraction strategy for individual workflows. This benchmarking analysis highlights a sensitive dual workflow capable of deciphering sex differences in heart.

### Conserved subcellular proteome of mouse heart

We next questioned the ability to sequentially resolve lipid and protein subcellular networks of the heart using our dual lipid-proteomic workflow and construct a molecular map of conserved subcellular molecular features. Here, we used a differential centrifugation-based subcellular fractionation workflow to resolve organelles of the heart^60^. Here, we obtained simultaneous heart subcellular lipidome and proteome profiling across 11 subcellular fractions (Fig 3a). For simplicity and improved quantitation of cardiac lipids including CL and PIP1 species, we employed hydro-BUME based lipid extraction and proteome profiling of all subcellular fractions. We quantified subcellular fractionation profiles for 7,026 protein groups (Supp Table 14). A protein marker list of 491 protein groups spanning 14 spatially separated membrane-bound and membrane-less subcellular niches was constructed based on their subcellular annotations across publicly available databases^65,70^ and prior reports^60,83^ with their data consistency across all biological replicates (Fig 3a, Supp Fig 5a-b, Supp Table 15). We trained subcellular niche-specific random forest (RF) classifiers using this conserved protein marker list and applied the trained RF models to predict protein localisation in each biological replicate. Overall, 3044 protein groups in the heart are spatially mapped to one primary subcellular niche across all biological replicates and with clear spatial resolution (Fig 3a, Fig 3b, Supp Fig3c, Supp Table 15). The median and mean RF probability scores for assigned proteins were 0.90 and 0.82, respectively, supporting high-confidence spatial assignment (Supp Fig 3d).

**Figure 3.**
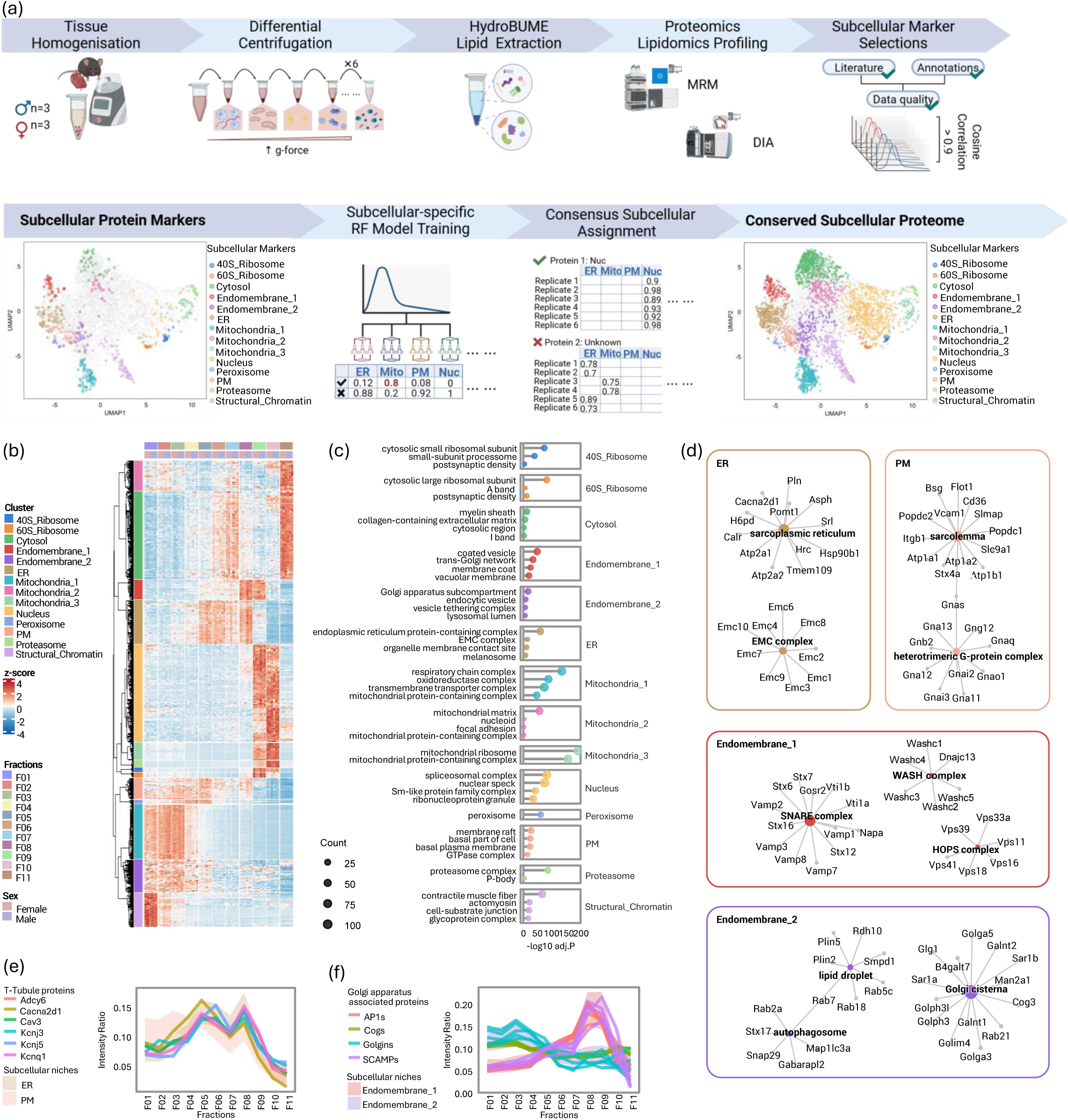
Spatial mapping of proteins defines the cardiac organellar proteome landscape. (a). Workflow of spatial classification of cardiac subcellular proteome. Left ventricle tissue samples derived from male and female hearts were subjected to homogenization and differential centrifugation-based subcellular fractionation, followed by Hydro-BUME lipid extraction where lipid extracts and protein extracts were processed and analysed via MS-based profiling. Subcellular protein markers were curated and visualised on UMAP with spatial distinctions. These proteins were used to train subcellular-specific random forest-based models to predict protein localisation probabilities across all subcellular niches. Replicate consensus filtering was enabled to preserve proteins with consistent localisation assignments as conserved cardiac subcellular proteome. (b) Heatmap of z-score normalised protein intensity ratios of the conserved cardiac subcellular proteome and their assigned subcellular niches from all subcellular fractions and different biological samples. (c). The top significantly enriched GOCC terms (Padj < 0.05) of proteins assigned to each subcellular niches. (d). Protein-term network plots of selected significantly enriched GOCC terms (Padj < 0.05) and the assigned proteins of ER, PM, Endomembrane_1 and Endomembrane_2 categories. (e-f). Profile plot of protein intensity ratio (0-1) against 11 subcellular fractions. The shaded areas represent the of 95% confidence intervals of fractionation profiles for proteins assigned to the subcellular niche of interest. Profiles of selected proteins (e) and protein complexes (f) are overlaid for comparison.

Functional enrichment analysis supported the predicted localisation of classified proteins (Fig. 3c). The spatial organisation observed in the current study showed strong agreement with our previous mouse heart subcellular proteome map, supporting reproducibility of the workflow in highly contractile tissue^60^. We highlight Endomembrane_1 and Endomembrane_2 clusters enriched with proteins associated with the Golgi apparatus and intracellular vesicular compartments, including endosomes, lysosomes and lipid droplets (Supp Table 16). Proteins annotated to endoplasmic reticulum, peroxisome, plasma membrane and proteasome were correspondingly enriched in ER, peroxisome, PM and proteasome clusters. Mitochondrial proteins segregated into three distinct clusters representing mitochondrial membrane (Mitochondria_1), mitochondria matrix (Mitochondria_2) and mitochondrial ribosomal (Mitochondria_3) compartments. Nucleus cluster was enriched with nucleoplasmic protein complexes such as splicesomal complex and histone methyltransferase complex.

Next, we focused on endomembrane-related clusters (ER, PM, Endomembrane_1 and Endomembrane_2) which exhibit distinct subcellular fractionation profiles (Fig 3d) and represent interconnected membrane compartments involved in essential trafficking and signalling events in striated muscles including cardiomyocytes. ER cluster contains ER-defining oligosaccharyltransferase and ER membrane protein complexes together with cardiomyocytes associated sarcoplasmic reticulum proteins (Hrc, Atp2a2, Atp2a1, Pln, Srl) (Fig. 3d). PM category was found with key components of cytoplasmic facing and plasma membrane-localising heterotrimer G-protein complex proteins as well as cardiomyocyte-characteristic sarcolemma proteins (Itgb1, Slc9a1, Atp1a1, Atp2b4) (Fig 3d). T-tubule is a specialised invagination of sarcolemma crucial for extra-intracellular actin potential transduction^23,84^. Consistent with this organisation, we identified canonical T-tubule proteins (Cacna2d1, Cav3 and Kcnj5) within the ER cluster, indicating co-fractionation with sarcoplasmic reticulum proteins and supporting the known spatial juxtaposition between T-tubules and the sarcoplasmic reticulum (Fig. 3e).

Further, Endomembrane_1 and Endomembrane_2 categories contain specialised vesicles and Golgi apparatus associated protein components but displayed distinct molecular compositions (Fig 3d). Endomembrane_1 category is found with vesicle tethering, sorting and trafficking machinery protein complexes including HOPS complex, SNARE complex and WASH complex proteins (Fig 3d, Supp Table 16). In contrast, Endomembrane_2 category contains vesicular proteins of lysosome, autophagosome and lipid droplets. Golgi apparatus-associated proteins are found in both Endomembrane categories but displayed distinctive molecular compositions. Proteins associated with trans-Golgi networks and intracellular trafficking associated proteins such as AP-1 adaptor complex proteins and SCAMPs proteins are found in Endomembrane_1, whereas Golgi stacks associated proteins such as golgin and Cog proteins are found in Endomembrane_2 (Fig 3e, Supp Table). This segregation resolved neighbouring yet distinct Golgi subcompartments across endomembrane clusters.

### Mapping subcellular lipidome with proteome

We then evaluated the performance of subcellular assignment analysis across protein and lipid levels. Here, following hydro-BUME-based lipid extraction, subcellular fractionation profiles of 842 lipid species of 54 lipid classes spanning 5 major lipid domains was performed, including glycerophospholipids, sphingolipids, fatty acids, prenol lipids and neutral lipids (including sterol lipids and glycerolipids) (Fig 4a, Supp Table 5). We subsequently assigned lipid fractionation profiles with distinct proteome-derived subcellular landscape, where the majority of lipids co-fractionate with membrane-bound subcellular niches including Mitochondria_1, ER, PM, Peroxisome, Endomembrane_1 and Endomembrane_2 (Fig 4a, Supp Fig 6a). This highlights the structural and membrane identity roles of lipids in organellar membrane organisation^45,47^. Interestingly, different lipid domains show spatially resolved patterns, inferring the differential subcellular organisation of different lipid domains (Fig 4a, Supp Fig 6a).

**Figure 4.**
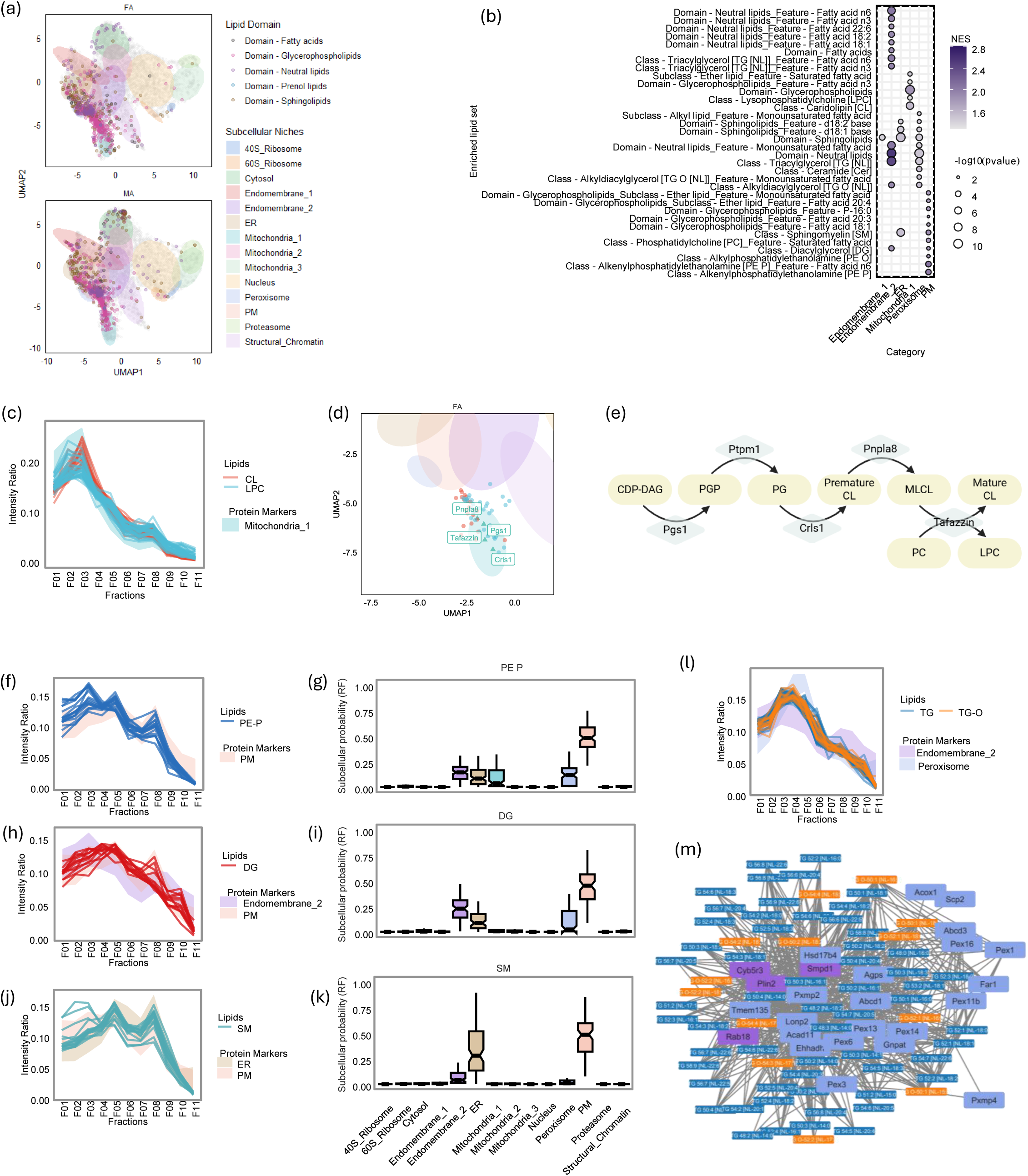
Spatial mapping of subcellular lipidome to proteome informs subcellular distribution of cellular lipids. (a). Representative UMAP plots of subcellular lipids data projecting to the UMAP embedding derived from subcellular proteome data. The shading area represents the ellipse of the conserved proteins of destinated subcellular category with confidence level of multi-variate t-distribution = 0.85. Individual lipids are coloured based their associated lipid domains. (b). LSEA based on the predicted localization probability of individual lipids across all investigated subcellular niches. (c), (f), (h), (j), (l). Profile plot of protein intensity ratio (0-1) against 11 subcellular fractions. The shaded areas represent the of 95% confidence intervals of fractionation profiles for proteins assigned to the subcellular niche of interest. Profiles of Mitochondria_1 enriched lipids (LPC, CL) (c), PM enriched lipids (PE-P) (f), Endomembrane_2 and PM enriched lipids (DG) (h), ER and PM enriched lipids (SM), (j) and Endomembrane_2 and Peroxisome enriched lipids (TG, TG-O) are overlayed with the protein-defining subcellular niches profiles for comparison. (d). Zoomed in UMAP plot of spatial distribution of CL, LPC lipids with cardiolipin-related biosynthesis and remodelling enzymes (Pgs1, Crls1, Pnpla8, Tafazzin). (e). Schematic illustration of proteins and lipids involved in cardiolipin biosynthesis ans remodelling. (g), (i), (k). Boxplot of subcellular localization probability distributions of PE-P (g), DG (i) and SM (k). (m). Protein-lipid interaction networks between TG and TG-O lipids with key lipid droplets and peroxisome-associated lipids. All interactions indicate the Pearson correlation > 0.9 between the connecting nodes.

To refine subcellular distribution of lipids from the heart, we applied the proteome-trained RF model to predict lipid localisation across 14 different subcellular niches (Supp Table 17). We retained only lipids with reproducible fractionation profiles across all independent biological replicates (n=6 hearts) (cosine correlation > 0.8) and determined mean subcellular localisation probabilities for each lipid species (Supp Table 18). Within each subcellular niche, lipids were ranked according to the niche-specific probability scores, and lipid set enrichment analysis (LSEA) performed to identify compartment-enriched lipid features (Fig. 4b, Supp Table 19). Here, cardiolipin (CL), a mitochondria inner membrane defining lipid^85–87^, was highly enriched in Mitochondria_1 niche (Fig. 4b-c), and lysophosphatidylcholine (LPC), a lipid product of cardiolipin remodelling^88,89^, displayed similar subcellular fractionation profile with CL (Fig. 4bc, e). Interestingly, proteins involved in cardiolipin synthesis (Pgs1, Crls1) and remodelling (Pnpla8, Tafazzin) also assigned to Mitochondria_1 are assigned in spatial arrangement with CL and LPC lipids, supporting a potential coordinated subcellular localisation of lipid metabolic pathways within mitochondria (Fig. 4d-e).

We observed plasmalogen phosphatidylethanolamine (PE-P) lipids, especially omega-6 fatty acid containing PE-P lipids associated with high PM localisation probabilities and subcellular fractionation profiles closely overlapping PM-associated proteins (Fig 4f-g). Diacylglycerol (DG) lipids are enriched in PM and Endomembrane_2 category, potentially indicating their subcellular distribution spans across plasma membrane and intracellular vesicles (Fig4 h-i). PE-P lipids are specialised lipids enriched in the inner leaflet of plasma membrane and lipid rafts that facilitate the biophysical property of membranes^90,91^, support signalling and scavenge reactive oxidative species. Sphingomyelin (SM) exhibited co-enrichment in ER and PM niches, with fractionation profiles and localisation probabilities distributed across these compartments (Fig. 4j-k). Previous studies have confirmed the enrichment of SM on plasma membrane, lipid rafts and specialised membrane invaginations such as caveolae and t-tubules in striated muscles^92–94^. Together with the observed co-fractionation of T-tubule proteins with ER-associated fractions, these results support SM potential localisation at sarcolemma and sarcoplasmic reticulum-adjacent T-tubules.

In addition, we report assignment in subcellular fractionation patterns of triacylglycerol (TG) and alkyl-diacylglycerol (TG-O) lipids in peroxisome and Endomembrane_2 (Fig 4l), known lipid classes enriched in lipid droplets and peroxisomes^95,96^. Here we reveal lipid droplet proteins and peroxisomal proteins that align in subcellular fractionation profiles (Pearson correlation > 0.9) with TG and TG-O lipids (Fig 4m). In the context of phosphoinositol (PI) lipids we observed distinctive subcellular fractionation patterns that share identical acyl chain compositions but equipped with differential phosphorylated inositol head groups (PIP1 38:4, PIP2 38:4) (Supp Fig6b). We predict using RF-based localisation both species primarily to the plasma membrane; however, PIP1 38:4 additionally spanned Endomembrane_2 and ER niches, whereas PIP2 38:4 showed higher ER-associated probability, indicating differential subcellular distribution associated with phosphorylation state.

In summary, we demonstrate the ability of using subcellular proteome knowledge to inform on the subcellular distribution of lipid species in the heart at a tissue level.

### Subcellular proteomics analysis reveals sex-dependent spatial distribution of proteins

To understand the proteome and lipidome subcellular distribution in a sex-dependent manner, we delineated the heart from male and female mice applied to this dual extraction workflow.

To identify protein and lipid species exhibiting sex-dependent subcellular patterns, we analysed their subcellular fractionation profiles using statistical models that report the level of confidence and magnitude of spatial differences. Here, we applied generalised additive models (GAMs) to model protein or lipid fractionation profiles without (null model) or with (alternative model) accounting biological sex as a categorial factor (Fig. 5a). These models at protein and lipid levels were compared through likelihood ratio testing where multiple-testing corrected p-values were reported, and to indicate the level of confidence in subcellular fractionation profiles between biological sex. To quantify the magnitude of spatial divergence, we calculated the mean pairwise cosine distance (1 − cosine correlation) of subcellular fractionation profiles between male and female. For lipidome subcellular distribution, we did not detect significant differences of the hearts in a sex-dependent manner (Supp Table 20), suggesting the conservation in lipid species at a subcellular scale in the heart.

**Figure 5.**
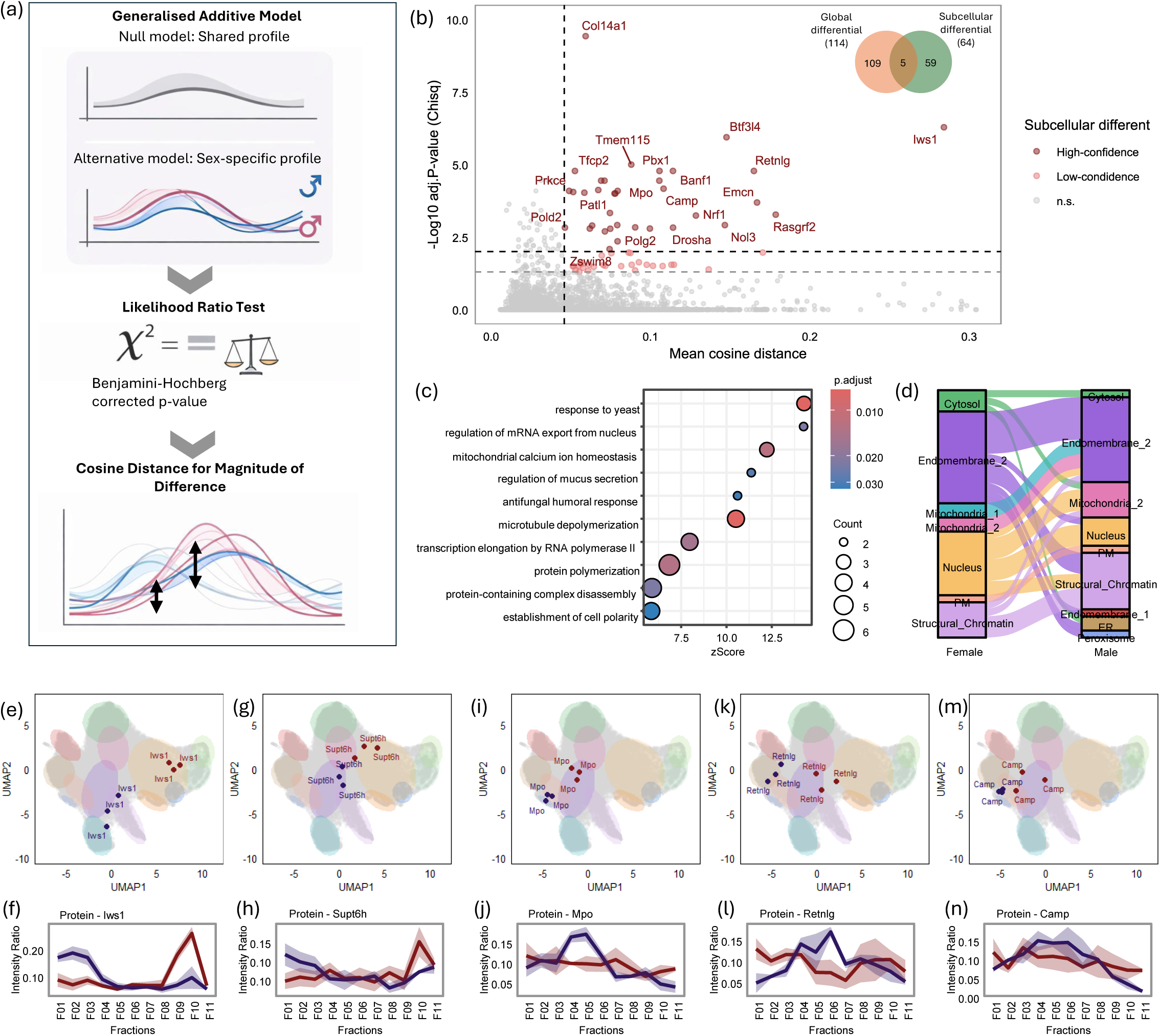
Sex-dependent differences in subcellular protein distribution. (a). Statistical workflow to identify proteins with sex-dependent subcellular protein distribution. Generalised additive models were fitted to protein fractionation profiles with (alternative model) or without (null model) biological sex accounted as a categorical factor. The two models were compared using likelihood ratio tests with BH-based p value correction. Pairwise cosine distance was calculated between sex-specific fractionation profiles to determine the magnitude of differences. (b). Scatter plot of 0-log10 transformed BH-corrected p value from likelihood ratio testing (Chisq) versus the mean cosine distance between female versus male-derived protein fractionation profiles. Proteins with Padj < 0.05 and mean cosine distance above 75^th^ percentile of entire protein cosine distance distribution are deemed significant with low confidence. Proteins with Padj < 0.01 and mean cosine distance above 75^th^ percentile of entire protein cosine distance distribution are deemed significant with high confidence. Venn diagram on the top right corner indicate the number of proteins with significantly different sex-dependent expression versus sex-dependent subcellular distributions. (c). Significantly enriched GOBP terms (Padj < 0.05) of proteins significantly different subcellular distribution between sexes. (d). Alluvial plot of proteins with differential subcellular distributions to their associated subcellular niches in female and male. (e),(g), (i), (k), (m). UMAP plot of the spatial arrangement of significantly subcellularly different proteins between male and female, including Iws1 (e), Supt6h (g), Mpo (i), Retnlg (k) and Camp (m). (f), (h), (j), (l), (n). Male (blue) and female (red) dependent fractionation profiles of significantly subcellularly different proteins, including Iws1 (f), Supt6h (h), Mpo (j), Retnlg (l) and Camp (n).

At protein level we identify sex-dependent subcellular distributions of 64 proteins (P.adj < 0.05, mean cosine distance > Q_0.75_ (distribution of mean cosine distances)) (Fig. 5b). Using this workflow, Camp, Chil3, Nol3, Myo9b, Fdx2 exhibit differential abundance together with distinct spatial patterns between sexes (Fig. 5b, Supp Table 21), indicating that most reported subcellular different proteins are independent of protein expression differences. Functional enrichment analysis of these subcellularly divergent proteins revealed enrichment in biological function networks associated with of immune cell related response, cytoskeleton organisation (microtubule depolymerization, protein polymerization), ribonucleoprotein related terms (regulation of mRNA export from nucleus, transcription elongation by RNA polymerase II) and mitochondrial calcium ion homeostasis (Fig 5c, Supp Table 22). We highlight that proteins that differ based on subcellular distribution include networks associated with Endomembrane_2, Nucleus and Structural_Chromatin categories (Fig 5, Supp Table 21).

Notably, the transcription elongation factor Iws1 displayed a highly significant sex-dependent spatial difference in the heart (Fig 5b). In male, Iws1 localised predominantly with the Structural_Chromatin cluster, suggesting its association with chromatin or nuclear envelope compartments, while in female aligns with Nucleus, indicating a nucleoplasm localisation (Fig 5e-f, Supp Fig). Iws1 interaction partner Supt6h also was shown to exhibit that same trend in heart spatial localisation (Fig 5g-h, Supp Fig). The co-expression of Iws1 has been shown in chromatin-bound and chromatin-free state to promote differential biological functions in transcription regulation, histone methylation and mRNA export^97–99^. Further, several proteins related to neutrophil granules (Mpo, Retnlg, Camp) also display sex-dependent fractionation profiles (Fig5i-n, Supp Fig), resembling ER or peroxisome-associated profiles in male hearts, while conversely aligning with the Endomembrane_2 niche in female (Fig5i-n, Supp Fig 7).

## Discussion

Here, we introduce Dual LiPA, a sequential lipid–proteomic analytical pipeline designed to identify, map and integrate lipid and protein features in the heart. By combining lipid extraction, high-recovery proteomic sample preparation, differential centrifugation-based subcellular fractionation and machine learning-based localisation prediction, this workflow enables integrated analysis of cardiac lipid–protein organisation at both global and subcellular scale. We first benchmark this dual extraction workflow for integrated proteome-lipidome analyses to demonstrate its performance. We then applied Dual LiPA to resolve the proteome and lipidome diversity of the left ventricular organelle landscape, revealing coordinated molecular networks across defined subcellular niches. Together, our findings show that biological sex contributes to cardiac molecular organisation at both abundance and subcellular spatial levels.

We first systematic evaluated the extraction workflows for integrated cardiac lipidomics and proteomics. Previously, multi-omics extraction strategies such as biphasic extraction workflow SIMPLEX^100^ and beads-assisted workflow BAMM^101^ have demonstrated the feasibility of simultaneous recovery of proteins, lipids and metabolites from the same biological sample. However, biphasic extraction workflows require careful handling of organic and aqueous layers, and bead-assisted extraction approaches introduce additional handling steps that may limit compatibility with simplified or automated workflows. Therefore, we sought to assess whether a simpler BUME-based extraction strategy could be adapted for concurrent lipidomic and proteomic profiling of cardiac tissue due to its monophasic solvent feature and high compatibility with robotic system^63^. Our result shows that although each extraction workflow introduced modest physicochemical preferences, the extraction-derived proteomes remained highly comparable to baseline and preserved the major biological differences between male and female heart proteome. In the current study, Hydro-BUME, a modified BUME-based workflow, provided a balanced strategy for cardiac lipid-protein extraction, particularly by supporting detection of biologically important lipid classes such as cardiolipins and phosphoinositides, without substantially compromising the extraction of the rest of the lipid classes and proteome.

At the global tissue level, Dual LiPA revealed sex-dependent differences in both the cardiac proteome and lipidome. Proteomic analysis showed that female and male hearts differed across pathways related to metabolism, immune regulation and redox biology align with previous reports^35,41^. Female hearts showed stronger representation of antioxidant and detoxification-associated features, whereas male hearts showed greater enrichment of innate immune-associated proteins, including complement-related components, suggesting that male and female hearts may adopt distinct molecular strategies to maintain cardiac homeostasis. In addition to these broader proteome-level differences, we identified known and candidate X-chromosome escapee proteins in the cardiac proteome. X-chromosome escape genes are genes that avoid complete silencing during X-chromosome inactivation, resulting in sex-biased gene dosage and expression^41,102,103^. Their detection supports the idea that cardiac sexual dimorphism may involve genetic mechanisms in addition to circulating sex hormones, which is particularly relevant because X-linked genes and X-chromosome escape have been implicated in sex-dependent immune regulation^41,104^. The lipidomic differences further supported this sex-dependent molecular organisation. The enrichment of omega-3-containing lipid species in female hearts is consistent with previous reports that women show higher conversion of α-linolenic acid to long-chain omega-3 polyunsaturated fatty acids, potentially influenced by sex hormones estrogen^105–107^. Female hearts were also enriched for ether-linked lipids, particularly plasmalogens, which can act as a reactive oxygen species scavenger^78,80^. In contrast, male hearts showed greater triacylglycerol abundance and lower level of lipolysis enzymes. Integration of lipidomic and proteomic data therefore suggests that male and female hearts differ in coordinated metabolic organisation, with female hearts showing signatures of redox protection and lipid utilisation, and male hearts showing features associated with lipid storage and immune-associated regulation.

At the subcellular level, lipidomics and proteomics integration is particularly informative because lipids and proteins jointly define organelle structure and function. Lipids support signalling, metabolism and niche-specific membrane environments for protein anchorage, while proteins regulate lipid synthesis, remodelling, transport and degradation. This lipid and protein interdependence is particularly important in the heart. Cardiolipin-enriched inner mitochondrial membranes support the organisation and anchorage of respiratory complexes required to meet the high metabolic demand of cardiac tissue^108^. Cholesterol- and sphingolipid-rich microdomains support the localisation of L-type calcium channels at t-tubules to maintain calcium signalling^84^.

Recent correlation profiling-based approaches such as LOPIT^50^and DOM^52^ have enabled cell-wide and tissue-wide protein localisation by combining subcellular fractionation-based proteomics and machine learning classification. However, equivalent lipid localisation remains challenging because lipids are often shared across multiple membranes and lack robust organelle-specific marker sets. Lately, computational framework C-COMPASS address this by using spatial proteomic profiles as references to infer lipid localisation^59^. Building on this principle, Dual LiPA used the conserved cardiac subcellular proteome as a spatial reference to reconstruct lipid distributions across the left ventricular organelle landscape. Here, we identified coordinated lipid–protein organisation across cardiac subcellular niches, including mitochondrial enrichment of cardiolipins and co-fractionation of TG and ether-linked TG species with lipid droplet- and peroxisome-associated proteins. These findings suggest that cardiac lipid distributions are organised in relation to protein-defined organelle architecture, supporting Dual LiPA as a practical framework for integrated spatial lipidomics and proteomics in the heart.

Following subcellular lipidome and proteome integration, we next investigated sex-dependent subcellular differences of cardiac proteome and lipidome. At proteome level, the transcription elongation factor Iws1 and its binding partner Supt6h showed distinct spatial patterns between male and female samples, suggesting possible sex-dependent regulation of chromatin-associated transcriptional complexes. Previous studies have shown that phosphorylation of Iws1 by Akt signalling can influence its chromatin association, raising the possibility that sex differences in Akt pathway activity may contribute to the observed spatial differences^97,99^. In addition, several neutrophil granule proteins showed altered fractionation patterns between sexes, suggesting potential differences in immune cell composition or granule organisation within cardiac tissue^109,110^. These observations indicate that sex differences in cardiac biology may extend beyond expression levels to include regulation of protein localisation and spatial organisation within cells.

Interestingly, sex-dependent lipid localisation differences appeared more subtle than protein localisation differences under baseline physiological conditions. This may reflect several biological and technical factors, including the broad distribution of membrane lipids across multiple organelles, cell-type heterogeneity in whole cardiac tissue, overlapping fractionation profiles between membrane-rich compartments, and the limited number of lipid species that uniquely define individual organelles. Therefore, while sex-dependent lipid abundance differences were readily detected at plasma and tissue level^111–114^, organelle-level lipid distributions may be more conserved in homeostasis. Previous subcellular organelle lipidomics in activated macrophages showed that inflammatory stimulation can remodel organelle lipidomes in a compartment-specific manner, suggesting that stronger lipid relocalisation may emerge under perturbation or injury contexts ^115^.

Despite the insights provided by this integrated proteomics and lipidomic analysis, there are a few notable limitations with the current pipeline. First, subcellular fractionation from complex tissue provides limited spatial resolution due to the overlapping fractionation profiles from neighbouring organelles, multi-localising lipids and proteins, or cellular heterogeneity within cardiac tissue. Increasing fractionation depth^59^ or incorporating click-chemistry based cross-linking approach may increase the subcellular spatial resolution and gain molecular insights of protein-protein and protein-lipid interactions^116^. Additionally, unequal lipid abundance across fractions may bias localisation predictions toward lipid-rich compartments, highlighting the need for complementary approaches such as antibody-based pull down of targeted organelle or protein of interests to validate lipid-protein spatial associations. Encouragingly, emerging spatial proteomics and lipidomics technologies including filter-aided expansion proteomics^117^ and transmission-geometry MALDI-MS^118^ provide valuable complementary routes to validate and extend fractionation-based spatial multi-omics. By adding spatial context to biochemical fractionation profiles, these approaches can help refine subcellular protein and lipid localisation and contribute to a more complete molecular map of cardiac subcellular organisation.

Collectively, this study establishes Dual LiPA as a framework for integrated spatial lipidomics and proteomics analysis in the heart. By linking molecular abundance with subcellular localisation, our approach provides a systems-level view of how lipid metabolism, protein organisation and organelle function are coordinated in cardiac tissue. Our results highlight the importance of considering spatial molecular organisation when interpreting sex-dependent cardiac biology. We provide a foundation for future studies examining how lipid-protein networks are remodelled during cardiac stress, injury and disease, with potential to inform the design of organelle-targeted therapies in precision medicine.

## Acknowledgments

This work was supported by the National Heart Foundation of Australia (DG: Vanguard), NHMRC project grant (DG: #1139489, 1057741), Future Fund (DG: MRF1201805), and the Victorian Government’s Operational Infrastructure Support Program. HF is supported by an Australian Government Training Program (RTP) scholarship and recipient of Bright Sparks Foundation scholarships (Baker Heart and Diabetes Institute).

## Additional Information

Competing interests: None

## Author Contributions

HF contributed to project coordination and design, experimental design, subcellular fractionation, heart dissection, mass-spectrometry sample preparation/data generation/data analysis, bioinformatics analyses, preparation of figures, manuscript drafting, and provided intellectual input. AR contributed to project coordination, experimental design, provided intellectual input, and manuscript review. KH contributed to lipidomics sample generation and experimental design, manuscript review. SSE contributed to heart tissue dissection and collection and provided intellectual input. TD, AF, and PM contributed to lipidomics sample generation and experimental design. DG contributed to project development and coordination, experimental design, data interpretation, funding acquisition, and manuscript drafting and review. All authors discussed the results and commented on the manuscript.

