## Supplementary Data (Figures S1-7) for "Dual-Lipa: A sequential multi-omic subcellular landscape map of mouse heart"

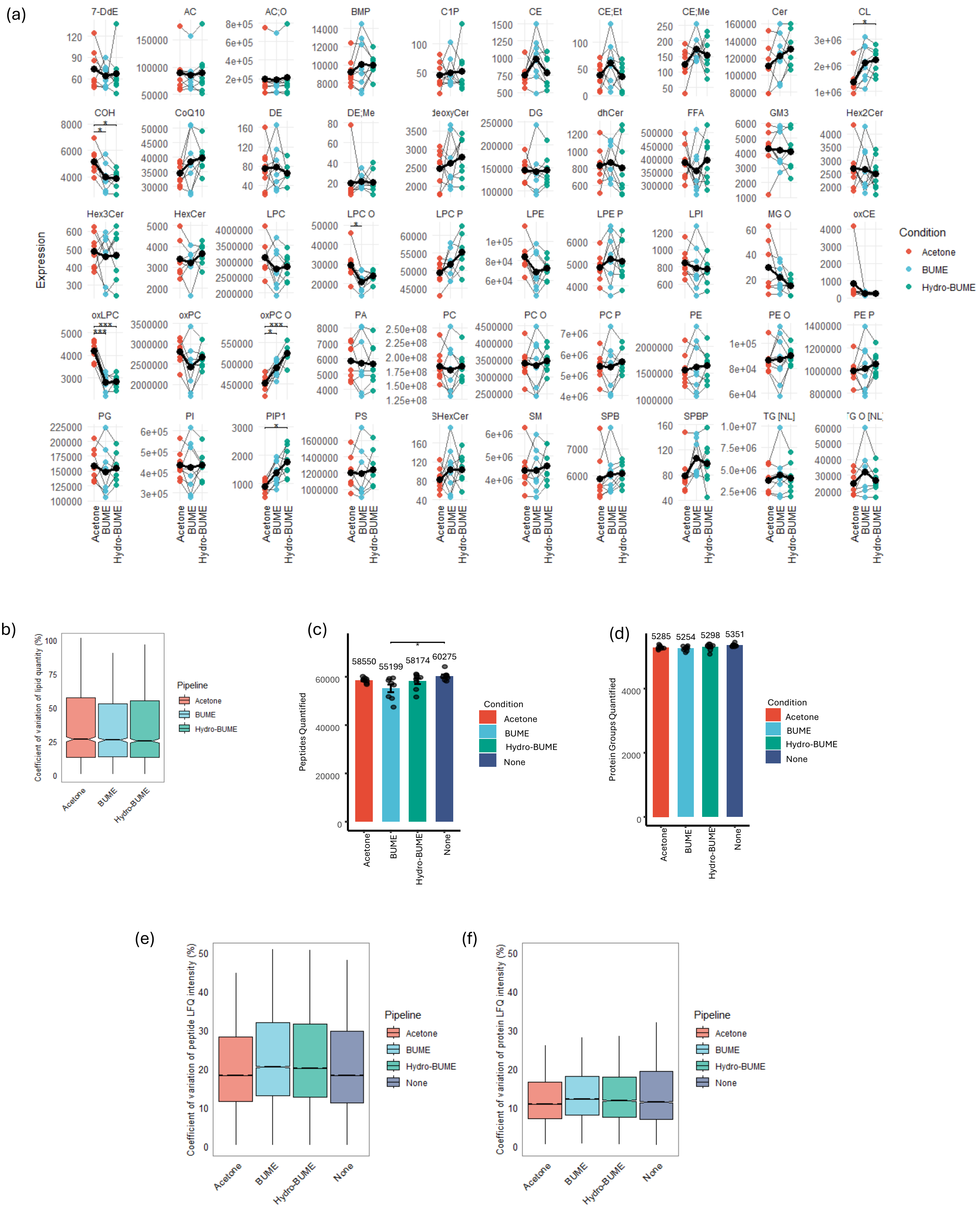

Supp Fig 1

Line plot of lipid intensities of induvial lipid class from the three lipid extraction pipelines. Lipid intensities was were using repeated-measures ANOVA with biological sample as the repeated measure, followed by Tukey-adjusted post hoc comparisons. Each points represent individual samples; asterisks indicate adjusted P values (*: Padj < 0.05, **: Padj < 0.01, ***: Padj < 0.001 ). (b). Boxplot of coefficient of variations (CV) of quantified lipid abundance (individual lipid species) from three extraction pipelines. Number of quantified peptides (c) and protein groups (d) from different extraction pipelines. Peptide and protein group numbers were analysed using repeated-measures ANOVA with biological sample as the repeated measure, followed by Tukey-adjusted post hoc comparisons. Bars show mean ± s.d.; points represent individual samples; asterisks indicate adjusted P values (Padj < 0.05). Box plot of CV of peptide (e) and protein group intensities (f).

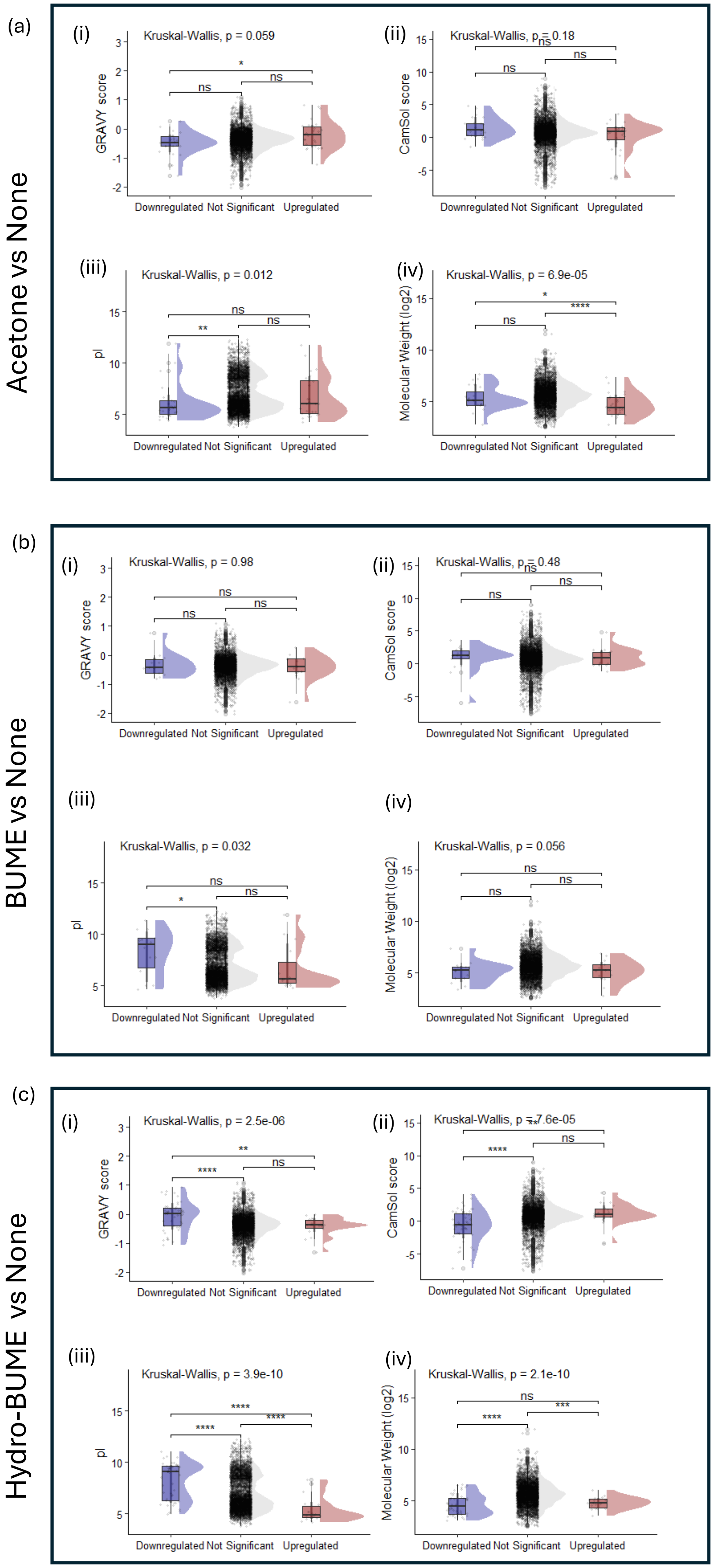
(a)
Acetone vs None
(i)
Kruskal-Wallis, p = 0.059
GRAVY score
Downregulated Not Significant Upregulated
(ii)
Kruskal-Wallis, p = 0.18
CamSol score
Downregulated Not Significant Upregulated
(iii)
Kruskal-Wallis, p = 0.012
pI
Downregulated Not Significant Upregulated
(iv)
Kruskal-Wallis, p = 6.9e-05
Molecular Weight (log2)
Downregulated Not Significant Upregulated
(b)
BUME vs None
(i)
Kruskal-Wallis, p = 0.98
GRAVY score
Downregulated Not Significant Upregulated
(ii)
Kruskal-Wallis, p = 0.48
CamSol score
Downregulated Not Significant Upregulated
(iii)
Kruskal-Wallis, p = 0.032
pI
Downregulated Not Significant Upregulated
(iv)
Kruskal-Wallis, p = 0.056
Molecular Weight (log2)
Downregulated Not Significant Upregulated
(c)
Hydro-BUME vs None
(i)
Kruskal-Wallis, p = 2.5e-06
GRAVY score
Downregulated Not Significant Upregulated
(ii)
Kruskal-Wallis, p = 7.6e-05
CamSol score
Downregulated Not Significant Upregulated
(iii)
Kruskal-Wallis, p = 3.9e-10
pI
Downregulated Not Significant Upregulated
(iv)
Kruskal-Wallis, p = 2.1e-10
Molecular Weight (log2)
Downregulated Not Significant Upregulated

Supp Fig 2

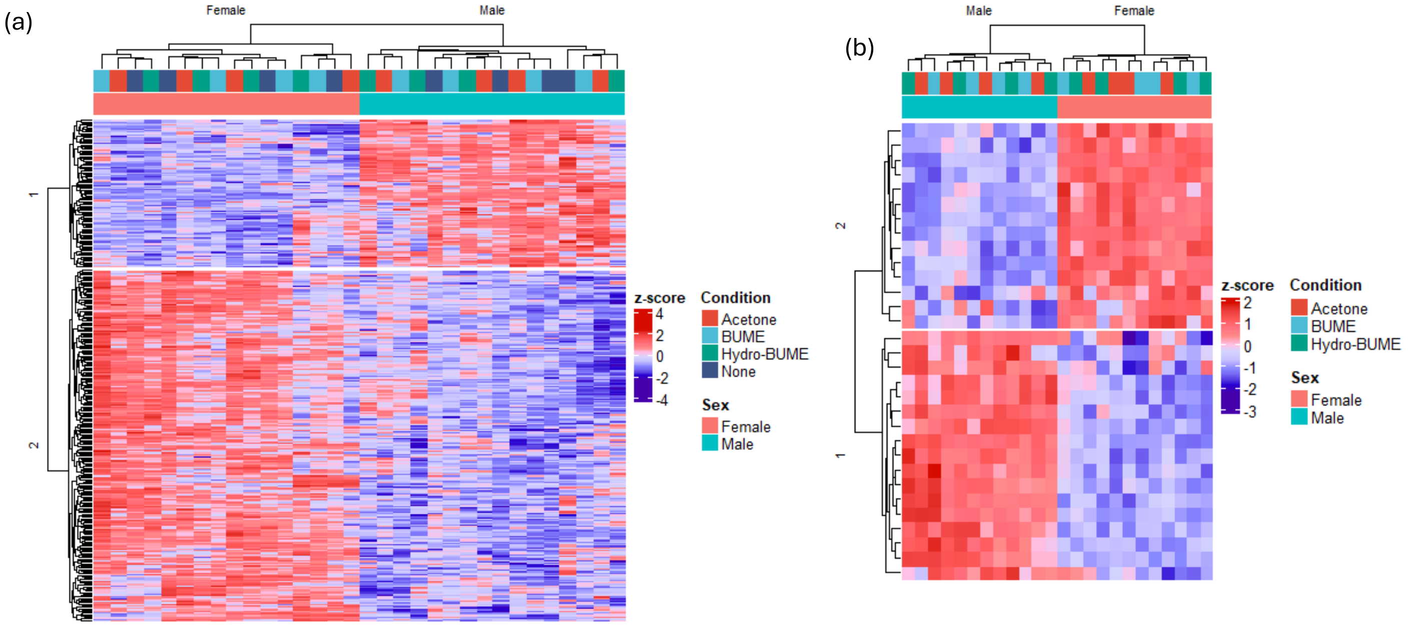


Supp Fig 3
Heatmap of significantly differential abundance proteins (a) and lipids (b) in male vs female left ventricles across all extraction pipelines.

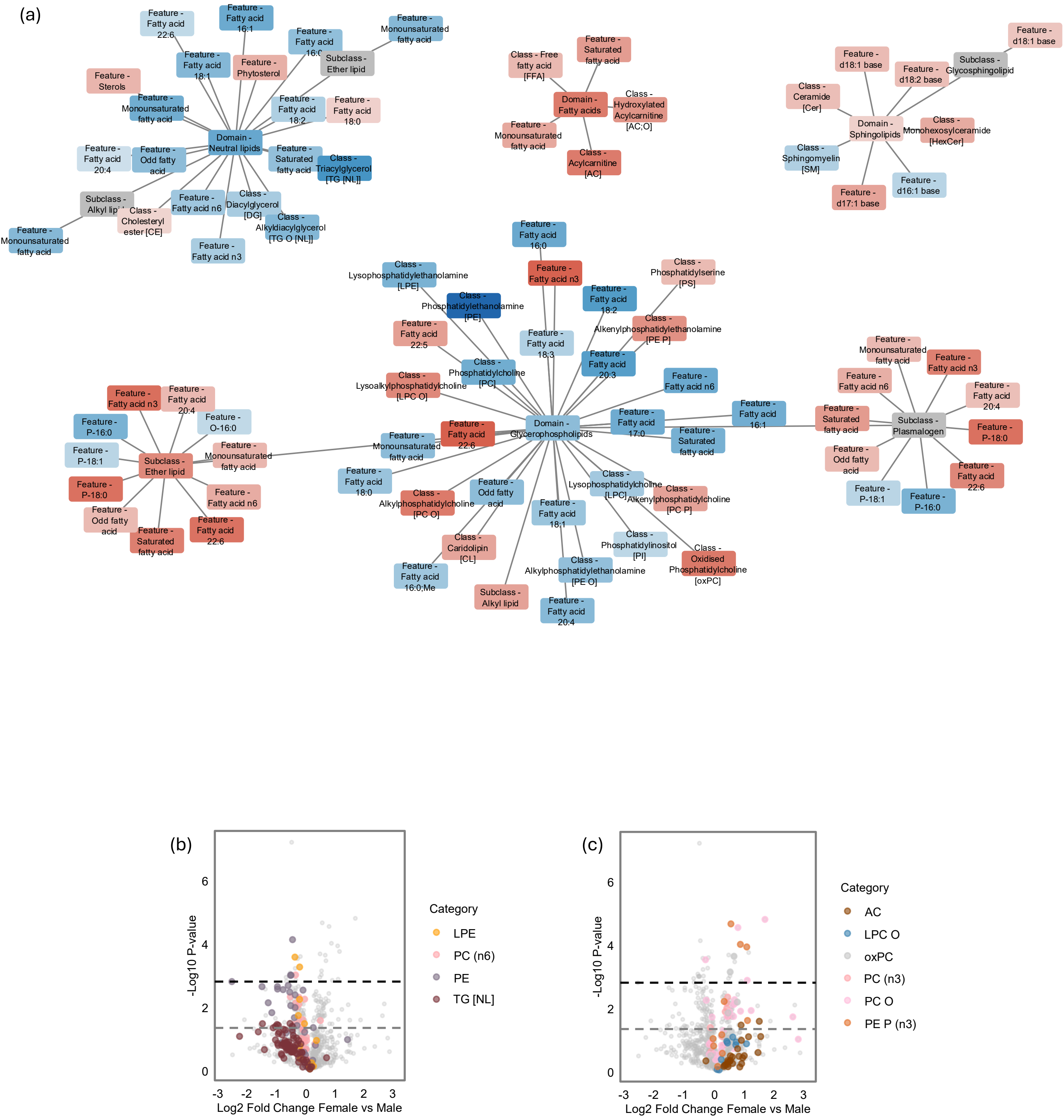


Supp Fig 4
(a) Network plot of Lipid domains, classes, subclasses and features eniched in male (blue) and female (red) left ventricles. Volcano plots of relative lipid abundance between male and female left ventricles with male (b) and female (c) enriched lipid classes highlighted.

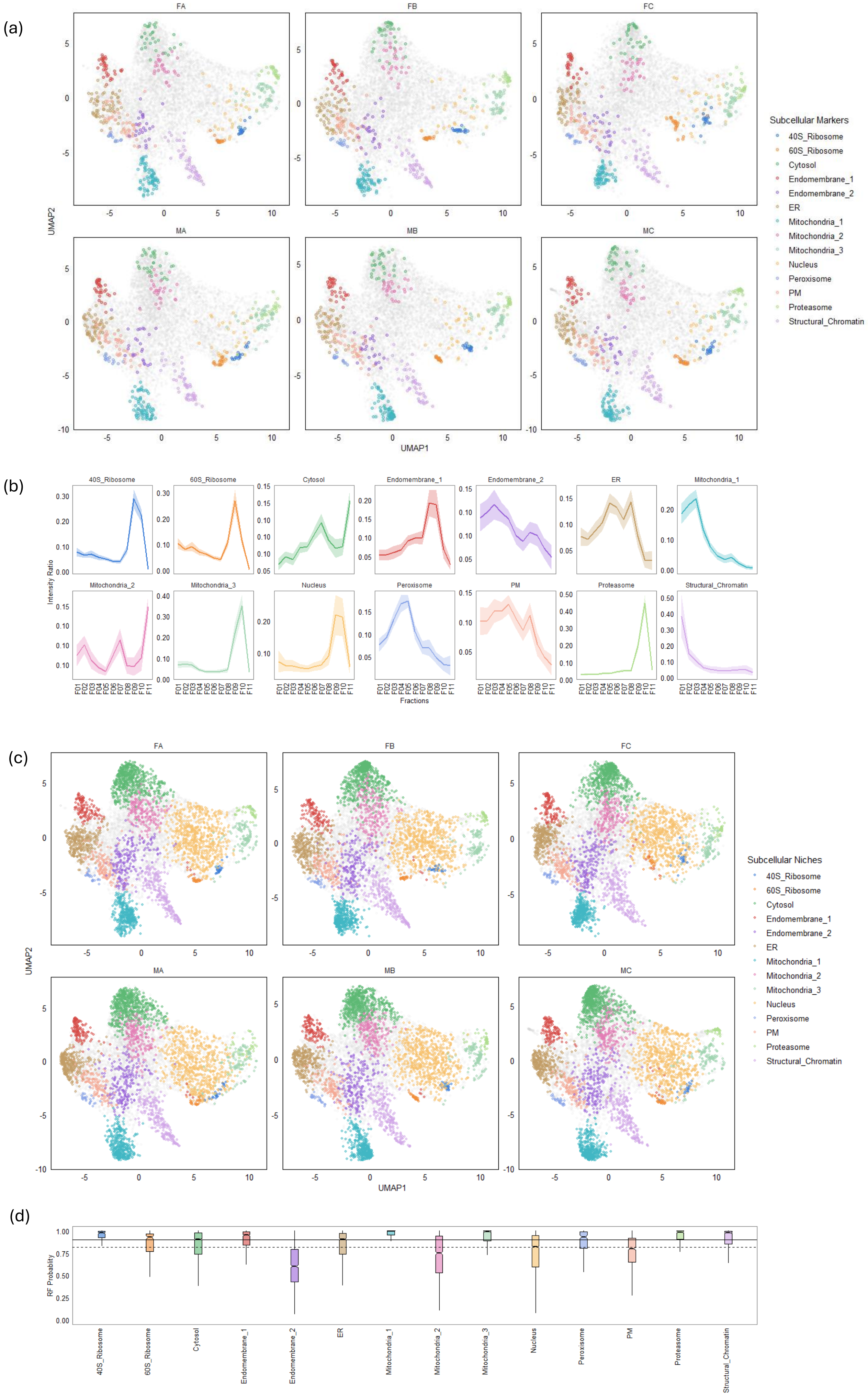
(a)
FA
FB
FC
MA
MB
MC
UMAP2
UMAP1
Subcellular Markers
40S_Ribosome
60S_Ribosome
Cytosol
Endomembrane_1
Endomembrane_2
ER
Mitochondria_1
Mitochondria_2
Mitochondria_3
Nucleus
Peroxisome
PM
Proteasome
Structural_Chromatin
(b)
40S_Ribosome
60S_Ribosome
Cytosol
Endomembrane_1
Endomembrane_2
ER
Mitochondria_1
Mitochondria_2
Mitochondria_3
Nucleus
Peroxisome
PM
Proteasome
Structural_Chromatin
Intensity Ratio
Fractions
F01 F02 F03 F04 F05 F06 F07 F08 F09 F10 F11
(c)
FA
FB
FC
MA
MB
MC
UMAP2
UMAP1
Subcellular Niches
40S_Ribosome
60S_Ribosome
Cytosol
Endomembrane_1
Endomembrane_2
ER
Mitochondria_1
Mitochondria_2
Mitochondria_3
Nucleus
Peroxisome
PM
Proteasome
Structural_Chromatin
(d)
RF Probability
1.00
0.75
0.50
0.25
0.00
40S_Ribosome
60S_Ribosome
Cytosol
Endomembrane_1
Endomembrane_2
ER
Mitochondria_1
Mitochondria_2
Mitochondria_3
Nucleus
Peroxisome
PM
Proteasome
Structural_Chromatin

Supp Fig 5

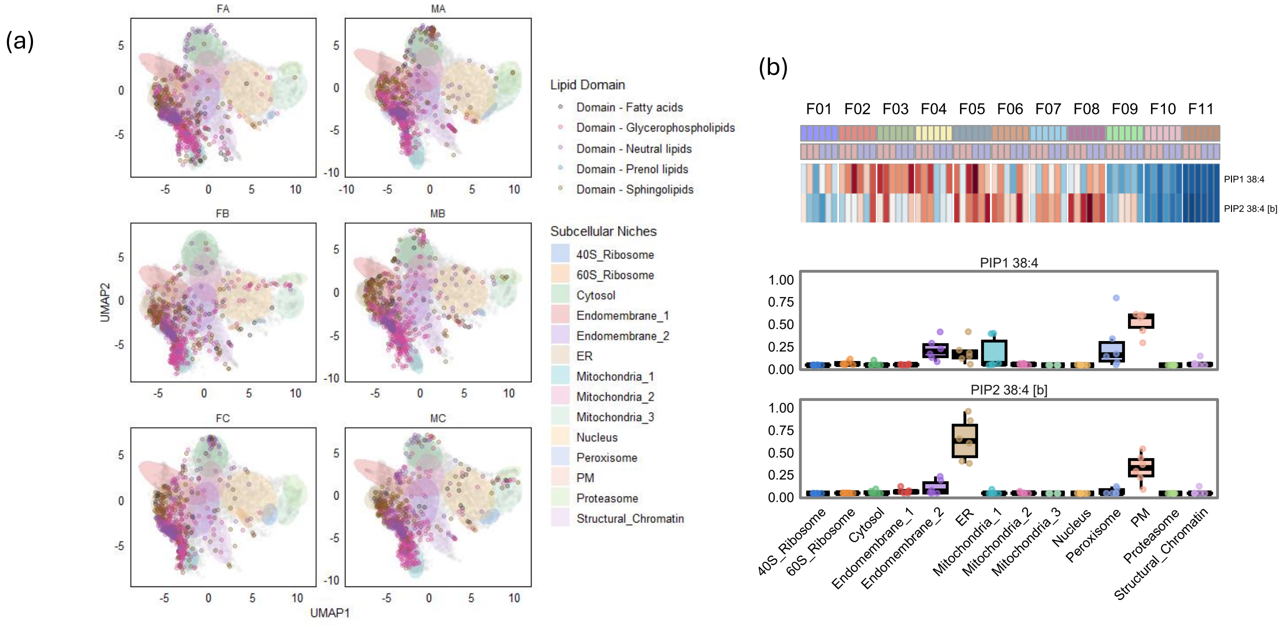


Supp Fig 6.

(a) UMAP plots of subcellular lipids data projecting to the UMAP embedding derived from each subcellular proteome data (F: Female, M: Male, A/B/C: biological samples). . The shading area represents the ellipse of the conserved proteins of destinated subcellular category with confidence level of multi-variate t-distribution = 0.85. Individual lipids are coloured based their associated lipid domains. (b). Heatmap of the mean intensity ratio of PIP1 38:4 and PIP2 38:4 [b] lipids from all biological samples. (c). Boxplot of subcellular localization probability distributions of PIP1 38:4 and PIP2 38:4 [b].

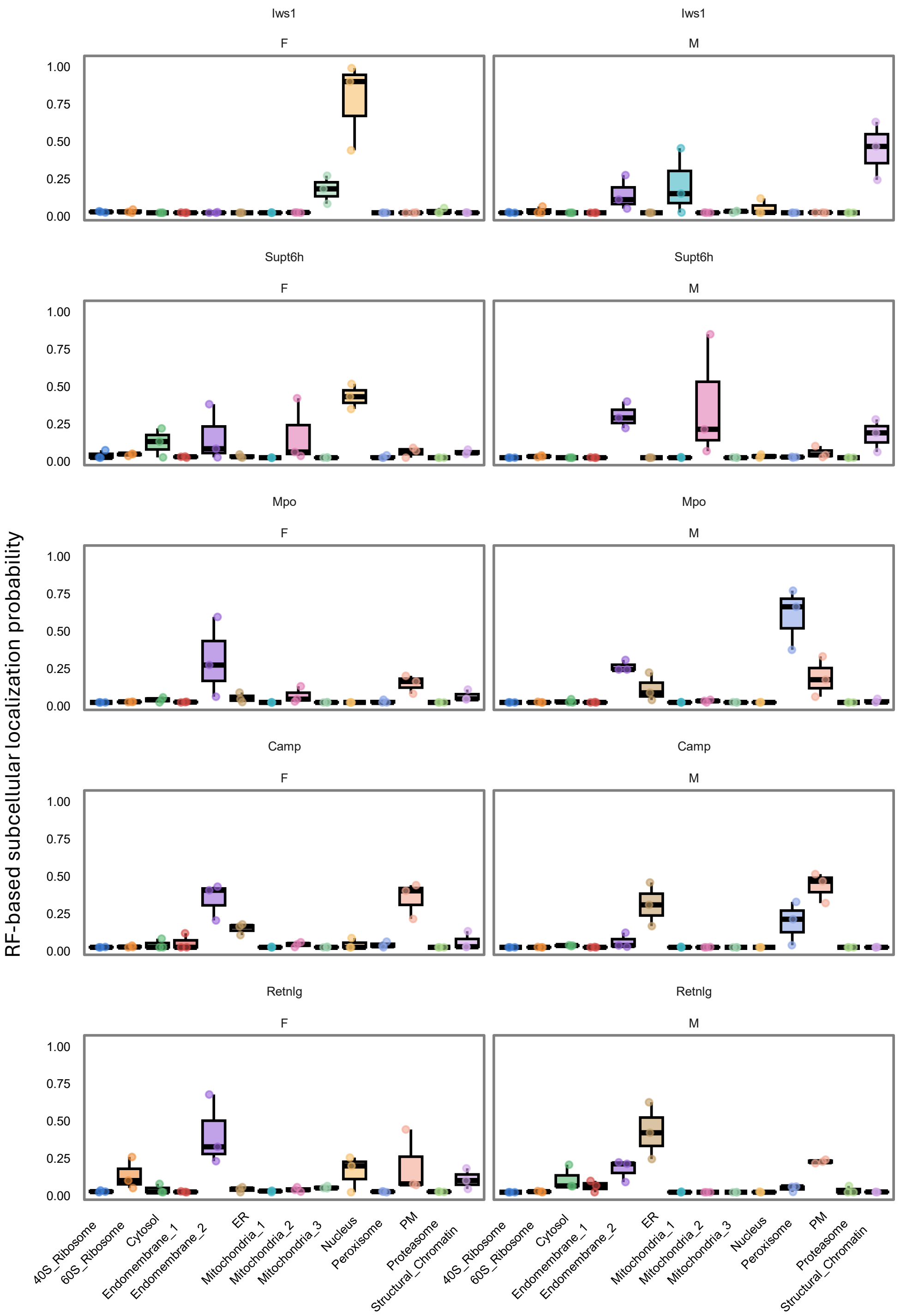


Supp Figure 7
Boxplot of subcellular localization probability distributions of Iws1, Supt6h, Mpo, Camp and Retnlg between male and female left ventricle subcellular proteome.
